# An outline on the chemical phenotype flexibility of forest species: an eco-metabolomics study of Pinus uncinata along an altitudinal gradient

**DOI:** 10.1101/2024.07.31.605974

**Authors:** Albert Rivas-Ubach, Ismael Aranda, Jordi Sardans, Yina Liu, María Díaz de Quijano, Ljiljana Paša-Tolić, Michal Oravec, Otmar Urban, Josep Peñuelas

## Abstract

The altitudinal distribution of plant populations is mainly determined by a set of environmental variables, including temperature, water availability, UV radiation, among others, which gradually shift with elevation. Therefore, altitudinal gradients in ecology could serve as “natural laboratories” providing insights into the phenotypic plasticity of natural plant populations. Plants can adjust their phenotypes to cope with specific environments. However, the adjustment capacity directly depends on the plasticity and flexibility of plant phenotypes. Plants growing at the edges of their distribution gradients may present limited flexibility due to the sub-optimal environmental conditions they experience.

We analyzed the foliar metabolomes of a mountain pine population in the Pyrenees to assess their chemical phenotypic flexibility along an altitudinal gradient. We found significant changes in foliar metabolomes across different altitudes, with the most contrasting foliar metabolomes observed at the lowest and highest altitudes. Trees growing at the boundaries of the altitudinal distribution considerably shifted their foliar metabolome compared to those at more central locations with an overall upregulation of sugars, amino acids, and antioxidants. Metabolomics analyses suggested higher oxidative activity at lower altitude, presumably due to the drier and warmer conditions. However, oxidative stress indicators were also detected at the tree line, potentially associated with chilling, UV, and tropospheric O_3_ exposure.

In addition to the inability of many species to keep pace with the rapid speed of climate change by migrating upward in altitude or latitude to find more optimum environments, their migration to higher elevations may be hindered by the presence of other environmental factors at high altitudes. Eco-metabolomics studies along environmental gradients can provide crucial insights into the chemical phenotypic flexibility of natural plant populations while providing pivotal clues regarding which plant metabolic pathway are prioritized to cope with specific environments.

## 1. Introduction

The geographical distribution range of a species is mostly constrained by a set of factors such as geography, biotic interactions, historical processes and environmental constraints. Environmental variables typically vary along gradients such as latitude and altitude (Körner, 2007), representing key factors delimiting the distribution of species populations. Latitudinal gradients may include large geographical features such as mountains, big rivers, or lakes that may abruptly interrupt the continuity of environmental conditions and the genetic flow within population individuals, especially in plants, which can drive the emergence of remarkable adaptation processes (i.e., emergence of subspecies). However, altitudinal gradients in mountain ranges typically present more gradual environmental changes over shorter geographic distances compared to latitudinal gradients. This reduces the probability of the emergence of particular varieties or subspecies with significant genetic modifications within the same population. In mountain regions, plant populations commonly present clear upper and lower distribution boundaries (Sexton et al., 2009). These limits are determined by a range of variables, including total solar radiation and UV light, water availability, temperature, and partial pressure of O_2_ and CO_2_ (Körner, 2007). Plants have the capacity to adjust different traits to cope with specific environmental conditions (Ghalambor et al., 2007). Therefore, the performance of plants located at the edges of the population altitudinal distribution could significantly differ from these located at intermediate altitudes that experience a more favorable environment (Fig. 1). In this context, the overall phenotypic plasticity of a plant species play critical roles in the acclimation and adaptation processes along environmental gradients, including altitude. The phenotypic plasticity refers to the capacity of a genotype to develop different phenotype configurations in response to the exposure to varying environmental conditions (Stager et al., 2024; West-Eberhard, 2005). However, considering that phenotypically plastic traits can exhibit varying responses to the environment, a distinction for those traits that are developmentally plastic (developmental plasticity) from the ones that are phenotypically flexible (phenotypic flexibility) was proposed (Piersma and Drent, 2003). While there is ongoing debate on whether these are distinct phenomena, most evidence indicates that developmental plasticity and phenotypic flexibility should be considered as separate aspects (Stager et al., 2024). On the one hand, developmental plasticity has been associated with the variability induced by the environment during the development of a single genotype. On the other hand, phenotypical flexibility refers to the reversible variations of an individual phenotype in response to the fluctuations of the environment (Piersma and Drent, 2003).

**Figure 1.**
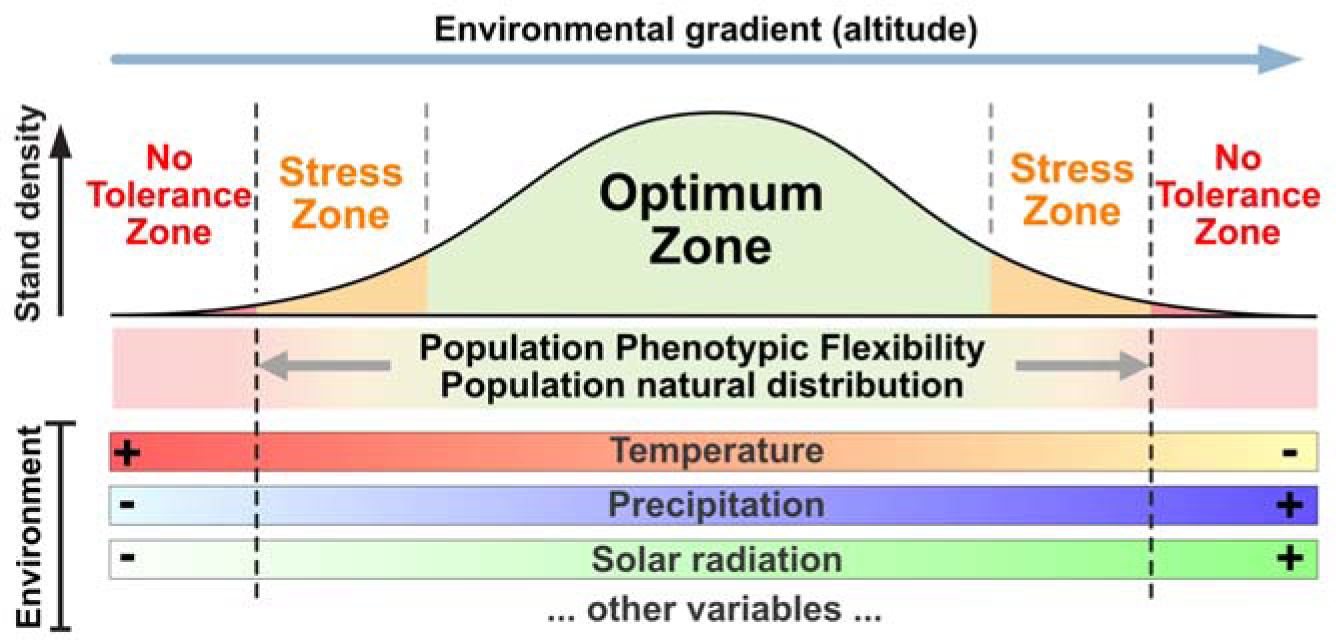
Conceptual framework of the phenotypic flexibility of plant populations along environmental gradients. Under a stable environment, plant populations would be distributed along environmental gradients (i.e. altitude) within the ranges comprising their phenotypic flexibility. This scenario would enable us to measure a good proxy of the boundaries of the phenotypic flexibility of natural plant population.

We hypothesize that organisms situated at the edges of the population’s natural distribution, where the environment is less favorable, would exhibit specific phenotypes nearing the limits of their phenotypic flexibility with a measurable degree of stress (Fig. 1). Therefore, altitudinal gradients can serve as excellent “natural laboratories” for understanding the impact of the environment on natural plant populations (Körner, 2007), especially in those individuals gorwing at the edges of the altitudinal distribution range (Schöb et al., 2013).

Studies have found clear relationships between altitudinal gradients with plant physiology (Cordell et al., 1999; Rajsnerova et al., 2015), ontogeny (le Roux et al., 2013), and even long-term plant genotypic selection at a local scale (Jump et al., 2006). Examples of described plant traits shifting with altitude include leaf mass area (Filella and Peñuelas, 1999; van de Weg et al., 2009), foliar nitrogen (N) concentration (Song et al., 2012), N-use efficiency (Cordell et al., 1999), phenological shifts (Meier et al. 2021), CO_2_ uptake capacity (Shi et al., 2006), and plant growth or overall fitness (Gottardini et al., 2016; Rajsnerova et al., 2015; Rossi et al., 2015).

Thus, those shifts would reflect the phenotypic flexibility of those specific tratis for the studied plant populations. Therefore, using altitudinal gradients is particularly advantageous in studies with anemophilous plant populations distributed along altitudinal gradients within relatively short linear distances between most extreme altitudes as pollination tends to be relatively uniform among individuals.

Changes in the physiology of plants are often a product of up-or downregulation of certain metabolic pathways (Brunetti et al., 2013). However, the overall metabolome of an organism experiences significant changes along environmental gradients rather than only specific pathways. A metabolome of an organism represents its entire set of low molecular weight (typically < 1000 Da) compounds (metabolites) present at a particular moment under specific circumstances. The metabolome comprises thousands of metabolites diverted in primary and secondary metabolic pathways of organisms (Fernie, 2007; Wang et al., 2019). Since those metabolites play crucial roles in organismal responses to environmental fluctuations it is being increasingly recognized the metabolome of organisms as their chemical phenotype (Fiehn, 2002). Indeed, extensive evidence demonstrates that plants quickly respond to environmental changes or stressors by modifying several aspects of their metabolomes (Sardans et al., 2011, 2020), which are commonly the first plant component responding to environmental fluctuations (Peñuelas and Sardans, 2009). Therefore, the different metabolome configurations of plants, which are constrained within their chemical phenotype flexibility, or hereafter referred to as metabolome flexibility, can directly reflect their functional status when exposed to particular situations. In this way, as a component of the overall phenotypical plasticity of a plant genotype, the metabolome flexibility which is predominantly shaped by climate and environmental gradients (Piersma and Drent, 2003), can significantly influence the capacity for acclimation and adaptation of the plant to new environments. We thus define the metabolome flexibility as the range of distinct metabolome configurations of an organism, which is constrained within specific thresholds, where it can survive and thrive. Metabolomic configurations beyond these specific thresholds defined by the metabolome flexibility would hinder the organisms’ success. Eco-metabolomic studies have shown large metabolome flexibility of plants under biotic and/or abiotic stressors (Bundy et al., 2008; Macedo, 2012; Rivas-Ubach et al., 2014; Robertson, 2005; Sardans et al., 2020, 2011). Nonetheless, the studies of metabolic responses of organisms to different stressors, or across individuals or species, are usually limited to a reduced pool of compounds (Aranda et al., 2018, 2017; Gargallo-Garriga et al., 2015; Rivas-Ubach et al., 2018, 2017, 2012; Sardans et al., 2020; Warren et al., 2011). In this way, eco-metabolomics research in altitudinal gradients not only can identify key molecular mechanisms involved in plant acclimation to different environments but can serve as a valuable tool to determine a reliable proxy of the metabolome flexibility of natural plant populations.

It is known that plants distributed on the lower and upper boundaries of the population altitudinal distribution may face the risk of experiencing events of forest decline and die-off under increased climate change conditions. Those plants, presenting metabolome configurations near the limits of their metabolome flexibility, could be forced to reconfigure their metabolomes into dysfunctional directions in order to cope with the new, more extreme environment. In fact, several studies have already shown increased forest die-off events in the most arid areas, typically coinciding with the lower distribution boundary of tree species, that has been linked to increased drought (Adams et al., 2017; Allen et al., 2010; Camarero et al., 2021; Gea-Izquierdo et al., 2021; McDowell et al., 2011; Pérez-Luque et al., 2021). Under scenarios of climate change, therefore, plant populations could face distinct outcomes depending on their metabolome flexibility (Ghalambor et al., 2007), competitiveness against other species, and surrounding topography, among other factors (Fig. 2). One of the major challenges plant species face is the rapid pace of climate change, which can hinder or impede their migration to higher altitudes and/or latitudes in search for more suitable environmental conditions (Huntley 1991). Selecting genotypes expressing better-adapted phenotypes for new climatic conditions may require several generations depending on the species and the existing genetic variability within the plant population (Rehfeldt et al., 2002). However, although natural plant populations may exhibit high levels of genetic variation, this variability is limited when confronted with the intense directional selection imposed by rapid climate change (Jump and Peñuelas 2005). Nevertheless, shifts in the distribution of natural plant populations in response to rising temperatures have already been documented (Walther et al., 2002), with upward altitudinal migrations (Grabherr et al., 1994; Lenoir et al., 2008; Liang et al., 2016; Peñuelas et al., 2007; Peñuelas and Boada, 2003). In this scenario, plant population would change their distribution range by shifting both lower and upper distribution boundaries to higher altitudes while maintaining the magnitude of their metabolome flexibility (Fig. 2a).

**Figure 2.**
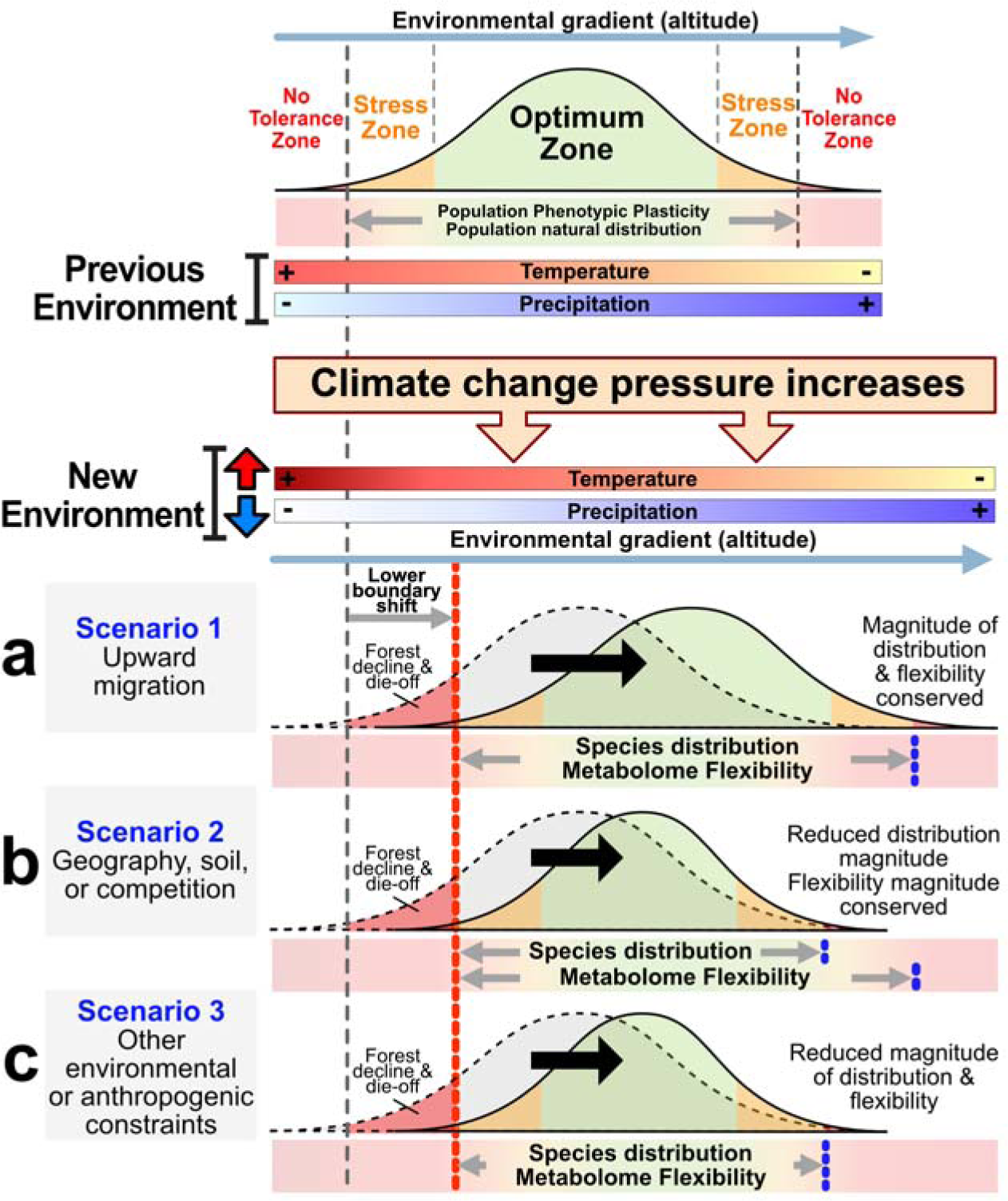
Conceptual framework illustrating three scenarios of the metabolome flexibility of natural plant populations along environmental gradients under the pressure of climate change. Scenario 1 (**a**) represents the upward migration of a plant population while conserving the magnitude of its distribution and the metabolome flexibility. Scenario 2 (**b**) represents a situation where a population cannot migrate to upper altitudes given the existence of physical constraints (i.e. top of a mountain; shallow soils) or competitors that impede the expansion of stands to higher altitudes. In this scenario, the potential expansion of the species distribution range has been reduced but keeps the same magnitude of the metabolome flexibility. Scenario 3 (**c**) suggests a situation where a plant population is blocked from moving upward given the appearance of new environmental constraints (i.e., wind, UV incidence, and tropospheric O_3_). In this scenario, the magnitude of the species distribution and the metabolome flexibility have been reduced.

However, plant populations in mountain ranges may also face different non-climatic obstacles (e.g., lack of soil lowering water holding capacity, presence of geographic barriers or competitors) blocking them to migrate to higher altitudes while still maintain the metabolome flexibility magnitude (Fig. 2b). Another potential situation would be the appearance or an increase of anthropogenic pressures (pollutants, tropospheric O_3_, grazing domestic livestock, etc.), or the intensity of other environmental factors at high altitudes such as UV incidence or wind or exposure that may hinder plant upward migration (Bais et al., 2018). In fact, an increased presence of pollutants such as O_3_ and N_2_O has been reported in high altitude areas of the Pyrenees in the Iberian Peninsula (Ribas and Peñuelas, 2006). Therefore, in this scenario, the distribution range and the magnitude of the metabolome flexibility of the population would be decreased (Fig. 2c).

Forest decline is a timely topic receiving special attention in the fields of ecology and ecophysiology (Choat et al., 2012; Galiano et al., 2011, 2010; Martínez-Vilalta et al., 2012; McDowell et al., 2022, 2011; Rowland et al., 2015; Wardle et al., 2004). As described with different tree species (Adams et al., 2017; Allen et al., 2010; Gea-Izquierdo et al., 2021; McDowell et al., 2011; Pérez-Luque et al., 2021), under the current trends of climate change scenarios, forest decline events are especially expected on the lower altitudinal distribution edge of plant populations, which typically coincide with the most xeric areas (Mátyás, 2010). However, this may not be the norm, and forest decline and die-off events could also occur in different locations along the altitudinal distribution of plant populations. The mountain pine (*Pinus uncinata* Ram.) is an autochthonous European tree species that dominates the subalpine forests (1400-2400 m a.s.l) in the Pyrenees (North-East Spain), representing the tree species that reaches the highest altitude in the mountain range. Some *P. uncinata* populations at the Pyrenees have exhibited increased mortality rates at their highest altitude areas of their distribution. Elevated mixing ratios of ozone (O_3_) have been identified as one of the main factors contributing to the events of forest decline and die-off (Díaz-de-Quijano et al., 2016, 2012). This phenomenon could indicate a potential scenario of enhanced stress for *P. uncinata* growing at high altitudes, which may impede their upward expansion in the current context of global warming (Fig. 2).

In this study, we characterized the metabolome of natural *P. uncinata* trees growing along an altitudinal gradient in the Pyrenees employing liquid chromatography coupled with mass spectrometry (LC-MS). Our primary goal is to explore and discuss the potential of eco-metabolomics approaches to elucidate the metabolome flexibility of plant populations and identify the main metabolic changes defining it. We discussed our results in the context of the existing literature, focusing on the roles of various key compounds in plant responses to the environment. As the ultimate response to the environment, this molecular-level information can play a crucial role in understanding the acclimation and adaptation processes of plant species in response to current climate change scenarios. Such insights are essential for assessing the future functionality of ecosystems.

## 2. Materials & Methods

### 2.1. Study site

The study was conducted in a population of *P. uncinata*along an altitudinal gradient in the central-eastern Pyrenees (Fig. A.1). The sampling collection was conducted in four different altitudes, i.e., Low (1,700 meters above sea level (m a.s.l.); 42°27’9”N - 1°47’32”E), Intermediate 1 (Int.1) (1,900 m a.s.l.; 42°27’22”N - 1°47’32”E), Intermediate 2 (Int2) (2,100 m a.s.l.; 42°27’51”N - 1°47’7”E), and High (2,300 m a.s.l.; 42°27’56”N - 1°46’42”E). Low and High altitudes represent the lowest and highest limits (tree line) of the natural distribution of *P. uncinata* in this area. The straight-line distance between Low-pines and High-pines is approximately 2.1 Km. Although the transport of viable pine pollen (up to 4.4% in Robledo-Arnuncio 2011) can still result in effective pollination over extensive distances, the proximity between pines of the same population across the four studied altitudes increases the likelihood of pollination among them. Consequently, this short-distance reduces the probability of substantial genetic variability within the same population. Pines growing at Low, Int.1, Int.2, and High are referred to as Low-pines, Int.1-pines, Int.2-pines, and High-pines. A summary of the environmental conditions for each altitude area is shown in Fig. 3a and described in detail in Díaz-de-Quijano et al., 2016 and 2012. Given that all environmental variables were significantly correlated, positively or negatively, with altitude (Fig. 3b), this study considers altitude as a multivariate representation of the local climate resulting from the combined effects of all environmental variables at each of the studied altitudes.

**Figure 3.**
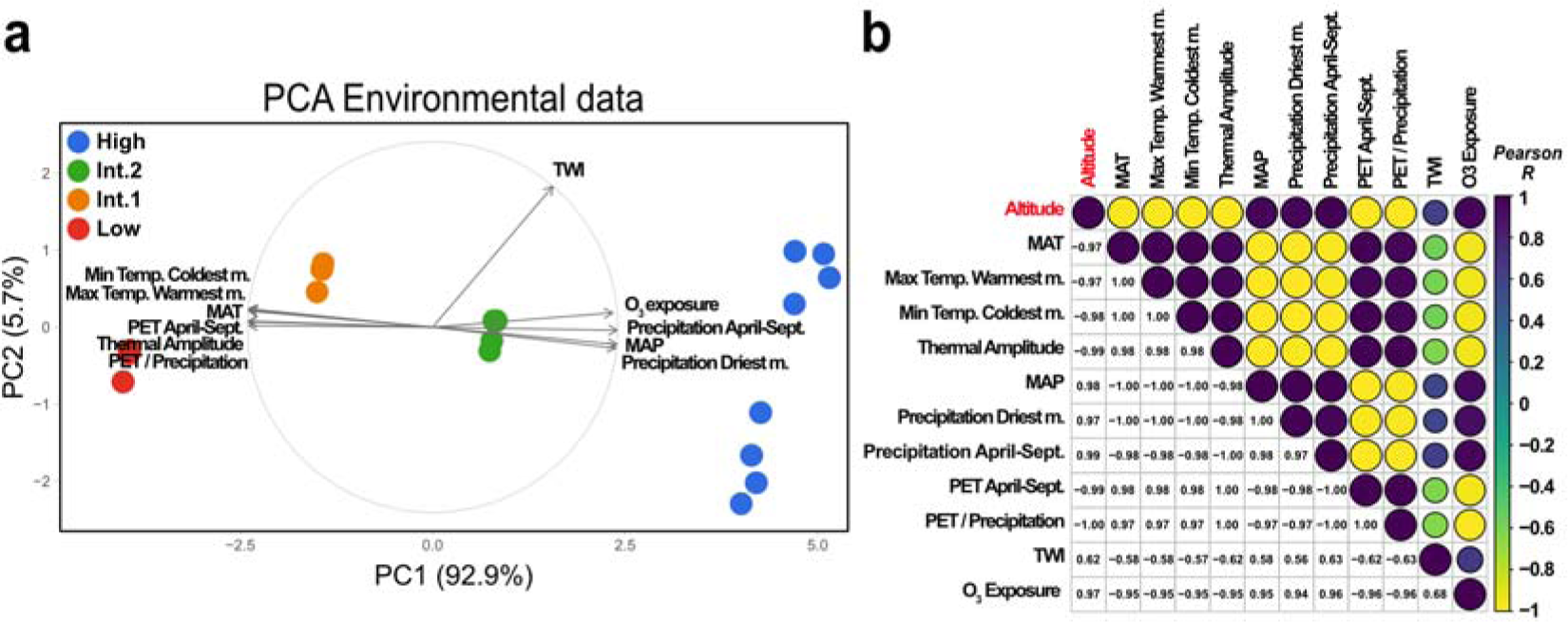
Summary of the environmental data along the studied altitudinal gradient. Biplot of the principal component (PC) 1 vs. PC2 of the principal component analysis (PCA) summarizing the environmental data changes across the different studied altitudes (**a**). Environmental data include mean annual temperature (MAT), maximum temperature warmest month, minimum temperature coldest month, thermal amplitude, mean annual precipitation (MAP), precipitation of the driest month, precipitation for April to September period, potential evapotranspiration (PET) for April to September period, PET/precipitation ratio, topographic wetness index (TWI) (Beven and Kirkby, 1979), and accumulated ozone exposure (see M&M) are shown for each altitude. Pearson correlations between all environmental variables and altitude are shown with an autocorrelation plot. Different colors and size of the circles indicate the strength of the correlation (Pearson correlation coefficient (R)), which range from dark blue (positive relationships) to yellow (negative relationships). Pearson r coefficients are also shown (**b**).

In addition, concentrations of O_3_ were monitored using passive samplers between 2004 and 2008 at all plots as previously described (Díaz-de-Quijano et al., 2009). This variable represents the long-term exposition of O_3_ in *P. uncinata,* and it does not represent the real O_3_ dose as estimated in other studies (Calatayud et al., 2016) but the exposition to tropospheric O_3_ independently on the diffusion rate into the foliar mesophyll.

### 2.2. Sample collection and processing

Ten adult individuals of *P. uncinata* per altitude were randomly selected as study subjects. All individuals were 5-6 m high with trunk diameters at breast height (DBH) of 25-35 cm. More details of studied pines can be found in Díaz-de-Quijano et al., 2016 and 2009. A small south-oriented sunlit branch was removed from each tree with a pole, and over 50 current-year needles were collected, packed in labeled Kraft envelopes, and immediately frozen in liquid nitrogen. All samples were collected in late summer (the second week of September 2011) when pine needles were already fully developed. Samples were collected between 11:00 – 14:30 local time on a sunny and calm day in order to diminish any effects of diurnal rhythms and changing meteorological conditions on plant metabolomes (Rivas-Ubach et al., 2013).

Frozen pine needles were processed in the laboratory following established eco-metabolomics protocols (Rivas-Ubach et al., 2013). Briefly, frozen needles were lyophilized for 72 hours at a constant pressure of 0.15 mbar and were subsequently ground in a ball mill at 1,600 rpm for 11 min. The resulting fine homogeneous powders were stored at −80 °C in polypropylene tubes until metabolite extraction.

### 2.3. Metabolite extraction for LC-MS

Metabolites were extracted with methanol:water (80:20) for a broad recovery of polar and semi-polar metabolites (t’Kindt et al., 2008). See *Metabolite extraction for LC-MS* section of Appendix A for detailed extraction procedures. The resulting extracts were stored at −80 °C until LC-MS analyses.

### 2.4. LC-MS analyses

LC-MS analyses were performed on a Dionex Ultimate 3000 HPLC (Thermo Fisher Scientific, USA) coupled to a high-resolution mass spectrometer (HRMS) Orbitrap XL equipped with a HESI II heated electrospray ionization source. The chromatography was conducted with a reversed-phase C18 Hypersil GOLD column (150 × 2.1 mm, 3 µm particle size; Thermo Fisher Scientific, Waltham, USA). The HRMS operated in FTMS (Fourier transform mass spectrometry) full-scan mode at a resolving power of 60,000. Full scan spectra were acquired over a mass range of 70–1000 m/z in both positive and negative modes. For each sample, 5 µL were injected in the instrument. Samples were randomly injected to distribute any alleged sensitivity variation in the instrument during the sequence. For more details on the LC analyses and HRMS acquisition parameters, see *LC-MS analyses* section of Appendix A and Rivas-Ubach et al., 2016c.

### 2.5. LC-MS chromatogram processing and dataset filtering

The LC-HRMS RAW files were processed with Mzmine 3.2.3 (Pluskal et al., 2010; Schmid et al., 2023). Briefly, the baselines of chromatograms were corrected, mass-to-charge ratios (m/z at MS1; exact masses) were detected, chromatograms for individual ions were generated with the ADAP chromatogram builder algorithm (Myers et al., 2017) and deconvoluted. Ions were thus aligned, and gap filled. Aligned ion chromatograms were manually checked in MZmine and clear noise features (e.g., m/z values shown across all chromatography) were removed. Metabolomic features (peaks with specific detected m/z and retention time (RT)) were matched against an in-house LC-MS metabolite library, containing the exact mass and RT values of over 600 primary and secondary plant metabolites. Compound identification using two orthogonal measurements (exact m/z and RT values) corresponds to a second level of compound identification confidence according to the metabolomics standards initiative (Sumner et al., 2007). Datasets were exported to CSV files (see Table A.1 for Mzmine parameters). The numerical values for each variable of the dataset corresponded to the relative abundance of the compounds and are suitable for relative comparative analyses (Gargallo-Garriga et al., 2014; Lee and Fiehn, 2013; Leiss et al., 2013; Mari et al., 2013; Rivas-Ubach et al., 2016b, 2014). Datasets were thus further filtered, normalized, centered and scaled before statistical analyses. The relative abundance of each metabolite feature was normalized by the total ion chromatogram. Then, metabolomic features were individually *mean-centered* and *auto-scaled* (divided by the standard deviation of the variable) (van den Berg et al., 2006) (see *LC-MS dataset filtering and normalization* section in Appendix A for details).

### 2.6. Statistical analyses

The metabolomics dataset was composed of one categorical variable (altitude) with four levels (Low, Int.1, Int.2, and High) and 10,391 metabolomic features (continuous variables) where 143 were identified (Table A.2). All statistical analyses were performed in R (R Core Team, 2021). Samples were first submitted to multivariate outlier tests based on the Mahalanobis distance for detecting potential outliers. Since no extreme outliers were detected when analyzing all altitudes together and no outliers were identified within each group, all samples were thus included in the subsequent statistical analyses. The outlier tests were performed with the *pca.outlier* function of the “mt” package (Lin 2024) (See Fig. A.2, Table A.3 and *Multivariate outlier tests* section of Appendix A and Table A.3 for details).

Permutational multivariate analysis of variance (PERMANOVA), performed with the Euclidean distance and 10,000 permutations, was used for detecting significant differences of the overall metabolome between pines growing at different altitudes. The analysis was performed for all altitudes together (Low vs. Int.1 vs. Int.2 vs. High) and between adjacent altitudes (Low vs. Int.1, Int.1 vs. Int.2, and Int.2 vs. High) (Table. 1). PERMANOVA was conducted with the function *adonis2* within “vegan” package (Oksanen et al., 2020). Individual oneway ANOVAs were additionally performed for each metabolomic feature (Table A.4). For a graphical representation of the pine overall metabolome differences across the four altitudes, the entire metabolomic fingerprints including all detected metabolic features were submitted to principal component analyses (PCAs). For plotting the PCAs, missing data of the metabolomics dataset were first imputed using the function *imputePCA* of the “missMDA” package (Husson and Josse, 2015; Le et al., 2008). Subsequently, PCAs were performed with the *PCA* function of “FactorMineR” package (Le et al., 2008). To assess the magnitude of metabolome changes across different altitudes, the Euclidian distances between all trees from different altitudes (fixed factor) were calculated as a proxy for metabolomic distances. The Euclidian distances were computed using the function *dist* of the “stats” package (R Core Team, 2021) and subsequently contrasted with one-way fixed effects analysis of variance (ANOVA) and HSD-post-hoc tests. ANOVAs were computed with the *aov* function of “stats” package” (R Core Team, 2021) and Tukey’s HSD pot-hoc tests were performed with the *HSD.test* function of the “agricolae” package (de Mendiburu, 2015).

Separate datasets of carbohydrates, amino acids, organic acids and other identified primary metabolites of interest, and phenolic compounds (compounds detailed in Table A.5) were generated from the entire dataset to summarize the main changes across different altitudes. Identified compounds were classified according to KEGG (Kyoto Encyclopedia of Genes and Genomes) and/or “Chebi” (Chemical Entities of Biological Interest). Categories “Flavonoids”, “Phenylpropanoids,” or “Phenols” were considered phenols. The datasets were submitted to PERMANOVA (Table A.6) and partial least square discriminant analysis (PLS-DA); a supervised clustering method that optimizes clustering of groups of samples (altitude) across two axes (components 1 and 2). PLS-DAs represent a summary of the contribution of metabolites to each of the altitudes across the 2D plane arising from components 1 and 2. PLS-DAs were computed with the *plsda* function included in “mixOmics” package (Rohart et al., 2017). Pathway analyses were conducted in Metaboanalyst 6.0 (Pang et al., 2024) to contrast High-pines vs. intermediate pines (Int.1 and Int.2 together) and Low-pines vs. intermediate pines. These analyses illustrate the main metabolic pathways impacted in each altitude (High or Low) in comparison to a central location of the population altitudinal distribution, which experience a more optimal environment. In addition, a principal component analysis (PCA) was performed for all compound classes separately and together to represent the overall variability across the different altitudes for each compound class. A correlation plot matrix based on Spearman correlation was generated for the phenolic compound dataset, including the variable altitude, to explore the phenolic compound correlation with altitude. Correlation plots were computed with the function *cor* of “stats” package (R Core Team, 2021) and plotted with *corrplot* of “corrplot” package (Wei and Simko, 2021).

According to the identified phenolics, the total relative abundance of phenolic compounds was estimated for each tree as follows:

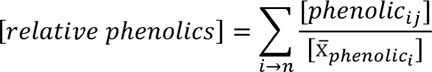

where *i* is a specific phenolic compound, *n* is the total number of phenolic compounds in the dataset, [*phenolic_ij_*] s is the relative abundance of phenolic *i* for the pine *j*, and 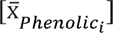 is the averaged abundance of the phenolic compound *i* across all 40 trees. Pairwise comparisons between altitudes using estimated marginal means were performed on the toral relative abundance of phenolic compounds to test for differences. Pairwise comparisons were performed with the *emmeans_test* function of the package “emmeans” (Lenth 2024).

## 3. Results

The PERMANOVA indicated that the overall metabolome composition of *P. uncinata* varied significantly across the four studied altitudes explaining up to 24% of the total metabolomic variability within the pine population (Table 1). Contrasts of consecutive altitudes also showed significant differences explaining the 14%, 13%, and 16% of the total variability for Low vs. Int.1, Int.1 vs. Int.2, and Int.2 vs. High altitudes, respectively.

**Table 1.** PERMANOVA tables of foliar metabolomes or *P. uncinata* across all altitudes and contrasting consecutive altitude pairs.

| All altitudes | Degrees of freedom | Sum of squares | R<sup>2</sup> | Pseudo-F | P |
| --- | --- | --- | --- | --- | --- |
| Altitude | 3 | 101,526 | 0.24 | 3.9 | <0.0001 |
| Residuals | 36 | 313,863 | 0.76 |  |  |
| Total | 39 | 415,389 | 1 |  |  |
| Low vs. Int.1 | Degrees of freedom | Sum of squares | R<sup>2</sup> | Pseudo-F | P |
| Altitude | 1 | 26,385 | 0.14 | 2.9 | <0.0001 |
| Residuals | 18 | 163,143 | 0.86 |  |  |
| Total | 19 | 189,528 | 1 |  |  |
| **Int.1 vs. Int.2** | Degrees of freedom | Sum of squares | R<sup>2</sup> | Pseudo-F | P |
| Altitude | 1 | 21,028 | 0.13 | 2.73 | <0.0001 |
| Residuals | 18 | 138,862 | 0.87 |  |  |
| Total | 19 | 159,889 | 1 |  |  |
| **Int.2 vs. High** | Degrees of freedom | Sum of squares | R<sup>2</sup> | Pseudo-F | P |
| Altitude | 1 | 30,666 | 0.16 | 3.66 | <0.0001 |
| Residuals | 18 | 150,720 | 0.83 |  |  |
| Total | 19 | 181,386 | 1 |  |  |

The PCAs of the foliar metabolome fingerprints showed clear clustering of the four altitudes across the first three principal components. PC1, explaining the 16.88% of the variability, clustered Low-pines, intermediate-pines, and High-pines (Fig. 4a). Int.1 and Int.2-pines showed the largest similarity between them with absolute overlap across the PC1 and PC2 (7,9%). However, the PC3 (7.37%) found clear clustering between Int.1 and Int.2-pines (Fig. 4b) in accordance with PERMANOVA results (Table 1), which accounts for 100% of the variance. One-way ANOVA on the metabolome Euclidean distances showed significant differences between altitudes (Fig. 4c). As expected, distances between individual pine metabolomes at the same altitude (*within-group* variation) showed the lowest metabolomics distance values except for Int.1 vs. Int.2 distance (*between-group* variation), which presented also low values if compared to the rest of distances. The Int.1-Int.1 and Int.2-Int.2 distances, and immediately followed by Int.1-Int.2 distance, showed the lowest values indicating higher metabolome similarity between pines growing at those altitudes. The metabolomic distance within Int.1 (*within-group variation*) was 15.6% smaller than the distance Int.1-Low; the two adjacent lowest altitudes of the study. Instead, the metabolomic distance within Int.2 was 20.3% smaller than the distance Int.2-High; the two adjacent highest altitudes of the study.

**Figure 4.**
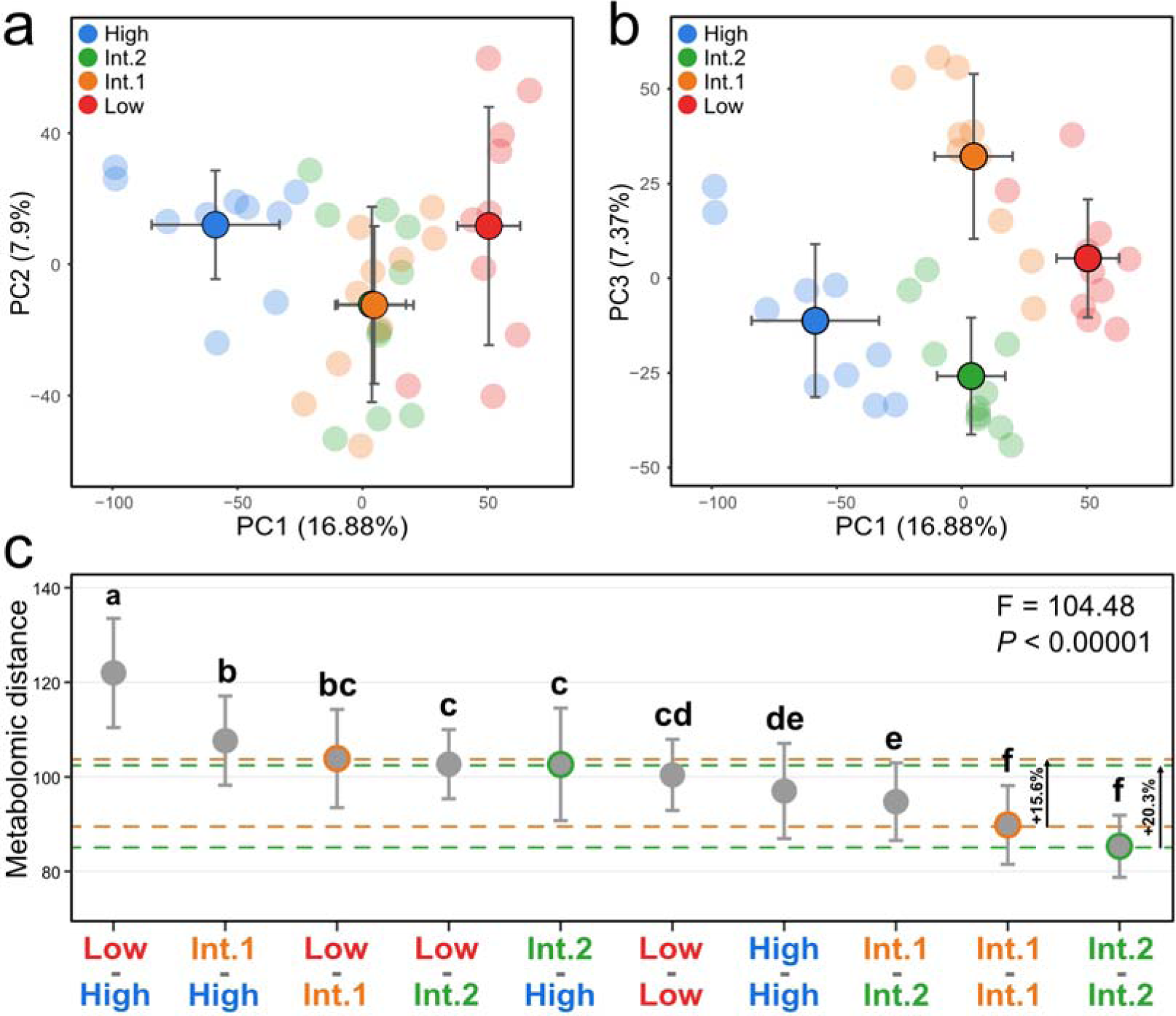
Metabolomic dissimilarity between the studied altitudes. Principal component (PC) 1 vs. PC2 (**a**) and PC1 vs. PC3 (**b**) of the score plot of the principal component analysis (PCA) performed with the metabolomic fingerprints of Pinus uncinata including all 10,651 detected features. Pines from different altitudes are shown with different colors (Low in red; Int.1 in orange; Int.2 in green; High in blue). The samples across both PCs of the PCAs are represented with faded dots and the average values for each altitude across each PC are shown in deep colored dots with black outlines. Standard deviation bars for the averaged values are shown for both axes. Euclidean distances (mean ± standard deviation), as a proxy of the metabolomic distance, between pine metabolomes of different altitudes (c). The horizontal orange dashed lines identify the Low-Int.1 and Int.1-Int.1 distances while the horizontal green dashed lines identify the High-Int.2 and Int.2-Int.2 distances. Different letters on the distances indicate statistical significance (P < 0.05) after one-way ANOVA and Tukey HSD post-hoc tests. Fisher’s F and P values of one-way ANOVA are shown.

The foliar profiles of carbohydrates, amino acids, and organic acids and other compounds of interest were all significantly different between pines growing at different altitudes (Table A.6; Fig. A.3). Even so, when performing PCA on all compound groups combined, we did not identify a specific group of compounds driving the differences by altitude (Fig. A.4). In general, the relative abundance of different identified carbohydrates varied across different altitudes although the relative abundances of most carbohydrates were found higher in Int.1-pines, followed by Low-pines, in comparison to High and Int.2-pines (Fig. 5a; Table A.4).

**Figure 5.**
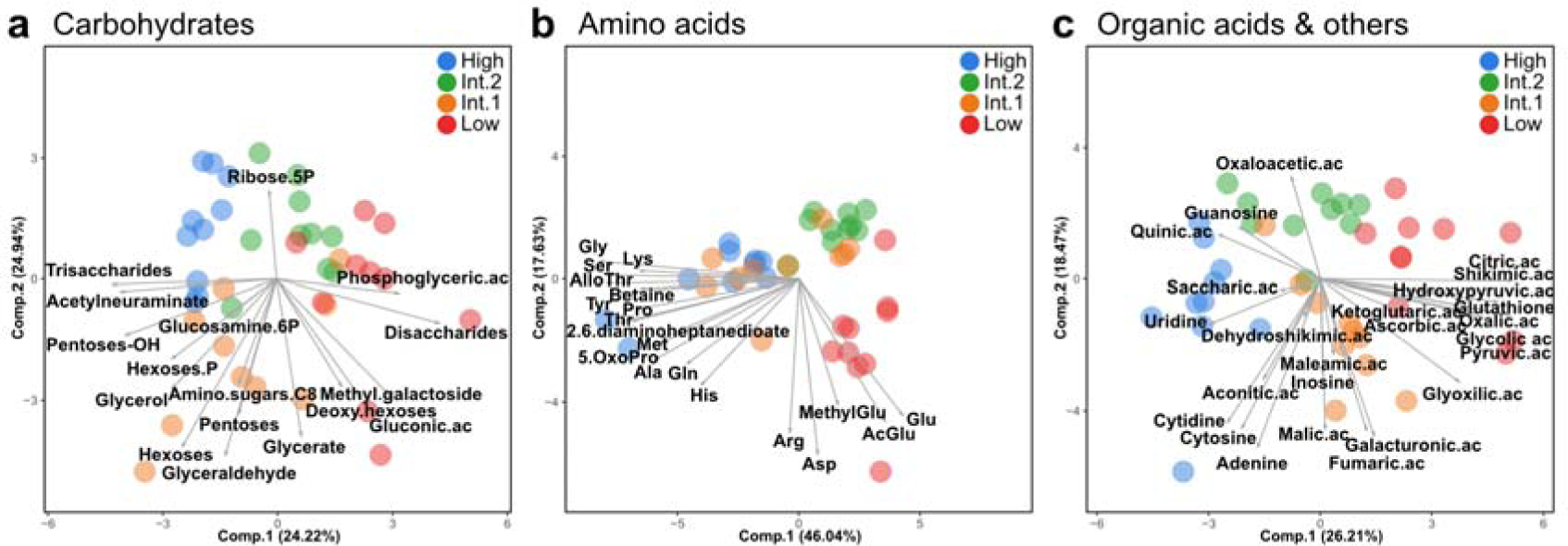
Primary metabolism shifts of P. uncinata along altitudinal gradients. Partial least square discriminant analysis (PLS-DA) (multivariate supervised ordination analyses) of the primary metabolism (carbohydrates, amino acids, organic acids and other metabolites of interest) of Pinus uncinata leaves. Biplots resulting from component (comp.) 1 vs. comp. 2 of the PLS-DA for carbohydrates (**a**), amino acids (**b**), and organic acids and other compounds of interest (**c**). Different altitudes are shown in different colors (Low in red; Int.1 in orange; Int.2 in green; High in blue). The loadings (identified metabolites) depicting the contribution of each variable (carbohydrates, amino acids, organic acids and others) to the altitude discrimination are shown as grey arrows in the PLS-DA plots.

Dissaccharides, deoxy-hexoses, and gluconic acid were significantly higher in Low-pines compared to the other of altitudes. Other carbohydrates such as trisaccharides and pentoses-alcohol, were found at higher relative abundance at High-pines. The PLS-DA and univariate analyses of amino acids revealed that the relative abundances of most identified amino acids were highest in High-pines, followed by Low-pines and Int.1-pines, with Int.2-pines having the lowest relative abundance across all amino acids (Fig. 5b; Table A.4). In general, High-pines presented higher relative abundances of glycine, glutamine, lysine, methionine, proline, serine, threonine, tyrosine, alanine, betaine, among others, while Low-pines presented higher concentrations of methyl-glutamic acid, acetyl-glutamic acid, glutamic acid, aspartic acid, and arginine (Table A.4). For the organic acid and other compounds of interest, PLS-DA and univariate analyses did not show a clear pattern across different altitudes. However, higher upregulation of several compounds such as glutathione and glycolic, glyoxylic, shikimic, citric, pyruvic, and hydroxypyruvic acids, was found at lower altitudes, especially in Low-pines (Fig. 5c; Table A.4).

Like the groups of compounds from the primary metabolism, the phenolic profiles showed significant changes across different altitudes (Fig. A.3; Table A.6). Most identified phenolic compounds changing significantly across distinct altitudes had larger relative abundances in High-pines, followed by Low and Int.1-pines (Fig. 6a; Table A.4). In a similar way as the amino acid profile, Int.2-pines had the lowest relative abundance of most phenolic compounds (Fig. 6a; Table A.4). This trend was also observed in the total relative abundance of phenolic compounds, with High-pines showing the highest values and Int.2 the lowest (Fig. 6b). Over 50% of the identified phenolic compounds significantly correlated with altitude (Fig. 6c). Compounds positively correlated with altitude included mangiferin, biochanin A, taxifolin, rhamnetin, fisetin, eriodictyol, ferulic acid, acacetin, 4-coumaric acid, 3-hydroxyphenylacetic acid, 2,4-dihydoxyacetophenone, and 3,2-hydroxyphenylpropanoic acid. Instead, eugenol, galangin, epigallocatechin, 3,4-dihydrobenzoic acid, and 2,3-dihydroxybenzoic acid correlated negatively with altitude.

**Figure 6.**
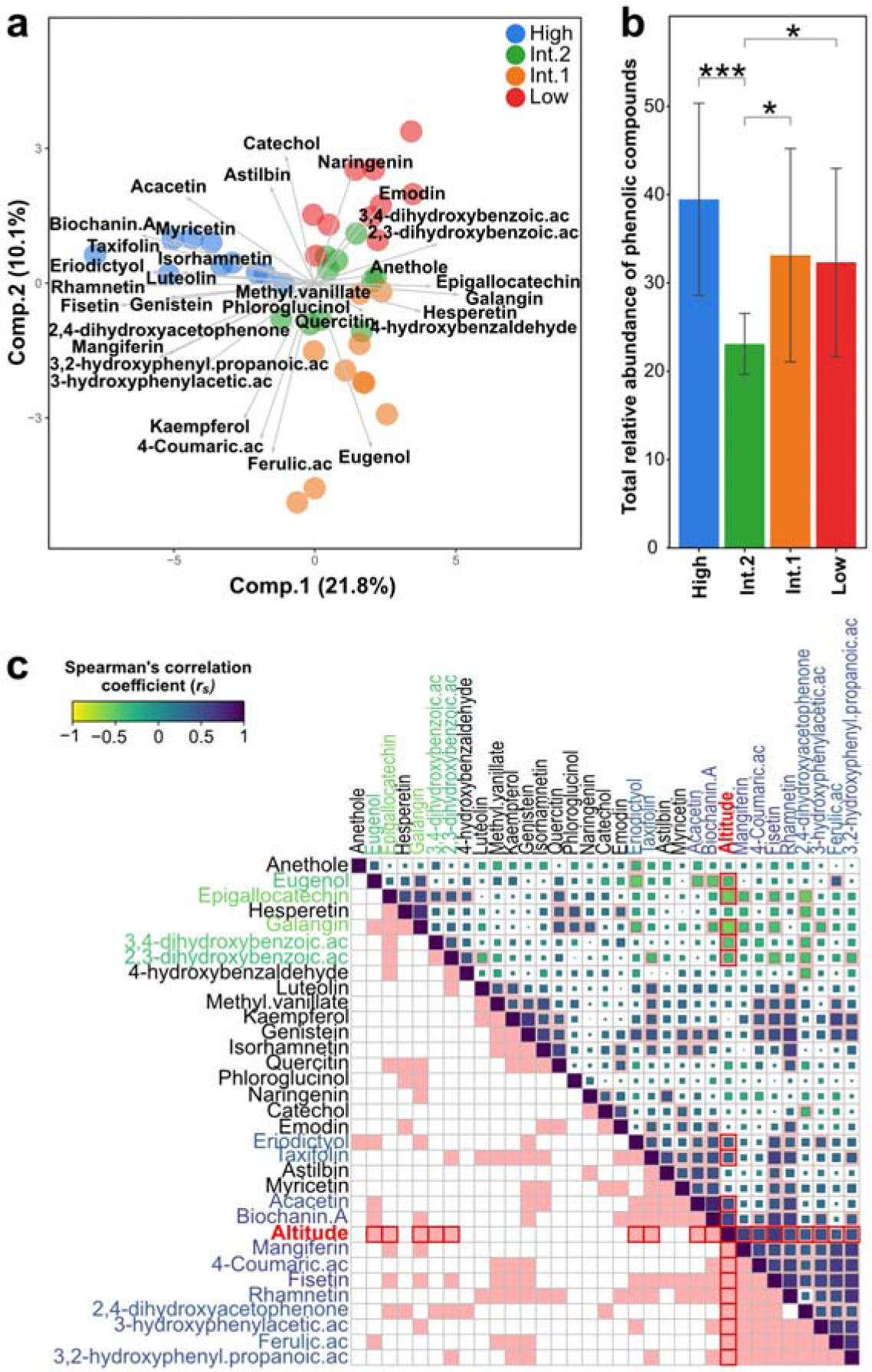
Phenolic profiles of P. uncinata along the altitudinal gradient. Biplot resulting from component 1 vs. component 2 of the PLS-DA for phenolic compounds (**a**). The loadings depicting the contribution of each phenolic compound variable to the altitude discrimination are shown as grey arrows. Relative abundance (mean ± standard deviation) of the total phenolic abundance for each of the altitudes (**b**). Asterisks denote statistical significance between different altitudes after pairwise comparisons of estimated marginal means (*** for P <0.001; * for P <0.05). Different altitudes are shown with different colors (Low in red; Int.1 in orange; Int.2 in green; High in blue) for the PLS-DA and the total relative abundance of phenolic compounds plot. Autocorrelation plot (Spearman correlation) between phenolic compounds across different altitudes (**c**). Altitude is included as an independent variable in the plot. Statistically significant correlations (P < 0.05) are shown in pink color while those significantly correlated with altitude are highlighted in red. Phenolic compounds significantly correlated with altitude are shown in different colors according to the Spearman correlation coefficient (scale of coefficients shown). Square size indicates the degree of correlation between variables with large squares for highly correlated variables and smaller squares for slightly correlated variables.

## 4. Discussion

### 4.1. Environmental metabolome flexibility of *P. uncinata*

As expected from the phenotypic flexibility of plant species, the overall metabolome of *P. uncinata* shifted with altitude (Table 1), which was directly related to a series of coordinated changes in environmental variables (Fig. 3). In accordance with the environment dissimilarity between those two altitudes (i.e., high temperature and evaporative demand at Low altitude, and low temperature at High altitude) (Fig. 3), PCA of the metabolome fingerprints of *P. uncinata* trees agrees with our hypothesis that plants growing at the edges of their natural distribution present the most divergent metabolome configurations as a response to less favorable environments (Fig. 4a, 4b). At the same time, the metabolomes of pines growing at central altitudes (Int.1 vs. Int.2) clustered together in the PCA performed with the PC1 vs. the PC2, indicating a more similar metabolome configuration between the intermediate altitudes if compared to High or Low-pines (Fig. 4a). Differences between intermediate altitudes clustered along PC3, which explained a similar proportion of total variability as PC2. These results suggest that metabolic changes in response to the environment intensify as trees approach the altitudinal edges of their natural distribution. Plants growing at the edges of their natural distribution range typically experience suboptimal environmental conditions, resulting in larger metabolome changes that may suggest a certain degree of stress (Fig. 1) (Sexton et al., 2009).The Euclidian distance, as a proxy of the metabolomic distance and includes the 100% of the metabolic variation, agreed with the PCA showed the High-Low distance as the largest metabolomic distance among all contrasted distances (Fig. 4c). The *within-group* metabolome variation of both Int.1 and Int.2-pines, obtained from the Euclidian distance between pines at those specific altitudes, got the lowest among all comparisons. However, these distances increased when intermediate pine metabolomes were contrasted against their adjacent pines growing at the altitudinal edges (*between-group variation*), showing an enhanced overall variation of approximately 15% for Low-Int.1 and 20% High-Int.2 distances. Therefore, this data obtained from eco-metabolomic measurements can serve as a multivariate proxy for assessing the metabolome flexibility of a plant population across their altitudinal distribution. In this regard, genetic variability can still be present across the distribution of a plant species, particularly along extensive latitudinal gradients (Ramírez-Valiente et al., 2010; Robledo-Arnuncio 2011; Robson et al., 2012; Sánchez-Gómez et al., 2013), thereby expanding the phenotypic plasticity thresholds of plant species. However, the metabolomic flexibility of plant populations - which involve the random collection a considerable number of samples from various sites - already account for the flexibility resulting from the genetic variability within the same population. Although more accurate values of the metabolome plasticity and flexibility of plant populations and species should be determined by combining measurements in natural and controlled environments with fitness assessments, this proxy of metabolomic flexibility can serve for contrasting plant populations from different locations, or different species, and shed light on acclimation processes in plant populations directly threatened by climate change. Our results clearly indicated that the environment shifts associated with a relatively small altitude difference (200 m constant between studied areas), as also detected with other phenotypic traits (Boscutti et al., 2018; Bresson et al., 2011; Miranda et al., 2021), significantly impact the overall metabolome configuration of trees, especially in those trees growing at the edges of the altitudinal distribution of the population. Thus, the magnitude of metabolomic changes can indirectly show the capacity of plants populations to reconfigure their metabolism in response to diverse environments, reflecting the inherent metabolome flexibility of the plant population.

Our results of the metabolome fingerprints of *P. uncinata* represent additional proof of the sensitivity of plant metabolomes to changing environments. Plants cope with environmental shifts and situations of stress require regulation of specific primary and secondary metabolic pathways (Bundy et al., 2008; Gargallo-Garriga et al., 2015; Rivas-Ubach et al., 2012, 2016; Sardans et al., 2020, 2011; Walker et al., 2022). We found significant changes in both primary and secondary metabolism of *P. uncinata* along the studied altitudinal gradient (Figs. 5 and 6; Table A.6). In addition, additional pathway analysis performed in Metaboanalyst (Pang et al., 2024) found distinct overall enriched pathways (pathways with a number of significantly changing metabolites) on pines growing at high and lower altitudes with respect both intermediate altitudes (Fig. A.5). Still, both altitudes exhibited few similarities in terms of impacted metabolic pathways, which can be directly linked to the gradual environmental shifts along the altitudinal gradient as the same metabolic pathway, according to the Kyoto Encyclopedia of Genes and Genomes (KEGG) maps, can be impacted in both altitudes but in a different manner (Figs. 5 and 6). These overall results of the metabolomic fingerprints provide key insights into the main metabolic components defining the metabolome flexibility of natural plant populations and, therefore, expose fundamental metabolic changes as key elements for acclimation and adaptation under scenarios of climate change.

### 4.2. Primary metabolism shift of *Pinus uncinata* along altitudinal gradients

Carbohydrates, in particular, non-structural carbohydrates (NSCs) that include starch and soluble sugars (glucose, fructose, sucrose, among others) represent a critical component of plant carbon storage. NSCs have been in the point of view in forest decline and die-off research, owing to their crucial osmoregulatory role that maintain turgor and lower plant water potential in periods of water stress (Blumstein et al., 2023; Dietze et al., 2014; Fernández-De-Uña et al., 2017; Gea-Izquierdo et al., 2023; Hartmann and Trumbore, 2016; Ingram and Bartels, 1996; Martínez-Vilalta et al., 2016). In general, high NSCs in plants have been associated to increased drought resistance (Garcia-Forner et al., 2016; O’Brien et al., 2014). However, NSCs include all soluble sugar species, thus failing to elucidate the mechanistic aspects of plant carbohydrate metabolism in response to water stress alone. Foliar sampling of *P. uncinata* was conducted at late summer, immediately after the driest and warmest period of the Mediterranean basin. Therefore, increased carbohydrate abundance can’t be directly related to carbon allocation for growth, as reported during the growing season in other studies (Rivas-Ubach et al., 2012). Carbohydrates in the studied trees were likely utilized to sustain cellular functions and to cope with environmental potential stressors (Ingram and Bartels, 1996; Keunen et al., 2013; Merchant et al., 2006; Rivas-Ubach et al., 2014; Rizhsky et al., 2004; Tauzin and Giardina, 2014; Warren et al., 2012). Monosaccharides hexoses (i.e., glucose, fructose, mannose), monosaccharides pentoses (i.e., ribulose, xylulose, xylose), disaccharides (i.e., sucrose, threhalose, maltose), trisaccharides (i.e., raffinose), glycerol and sugar alcohols are soluble carbohydrates generally found in higher abundances in plants exposed to water stress (Chen and Jiang, 2010; Irigoyen et al., 1992; Pamuru et al., 2021; Sánchez et al., 1998; Sardans et al., 2020). We observed a general trend of higher soluble carbohydrates abundance at the two lowest altitudes (Int.1 and Low), particularly disaccharides and monosaccharides hexoses, suggesting an increased osmotic function compared to higher altitude pines (Int.2 and High) (Fig. 5a; Table A.4). These metabolomics results align with the lower water potentials commonly observed in trees growing at the most xeric areas of their distribution (Jin et al., 2023). In addition, upregulation of gluconic acid, found at higher relative abundance in both Low-pines, and followed by Int.1, has been related an increased plant water stress as it may have the capacity to promote root growth and aquaporin activity (Han et al., 2019; Li et al., 2020). However, the function of gluconic acid in plants remains unclear and needs further mechanistic understanding (Han and Yuan, 2009; Sardans et al., 2020). Nevertheless, the pine carbohydrate overall response to an alleged water stress at the lowest altitudes do not follow a direct relationship with the environment as Int.1-pines showed the highest relative abundances of several compounds that did not change significantly with Low-pines or High-pines (Table A.4). The protection ability of soluble sugars in plants under oxidative stress has been typically connected with signaling triggering to the synthesis of reactive oxygen species (ROS) scavengers (Van Den Ende and Valluru, 2009). However, it has been proposed that sugars might also act as protective compounds against *in planta* against ROS, principally when found at high abundances (Keunen et al., 2013). Although the action mechanism is still unclear, high relative abundances of hexoses-phosphate, pentoses alcohol, and trisaccharides at High-trees could partly be associated with the increased foliar oxidative stress already reported on those pines (Díaz-de-Quijano et al., 2016, 2009). Different-omics analyses targeting on the dynamics of carbohydrate metabolism of plants under diverse stressors are necessary to obtain new mechanistic insigths into the potential of sugars acting as ROS scavengers.

Amino acids, and like many other metabolites, are often part of different primary and secondary metabolic pathways with important roles in plant regulation and stress signaling (Rai, 2002). Like carbohydrates, a higher abundance of amino acids has been related to plant developmental changes during the growing season (Gargallo-Garriga et al., 2014; Rivas-Ubach et al., 2014, 2012). However, beyond the growing season, several studies have associated overall upregulation of several amino acids with plant stress (Batista-Silva et al., 2019; Pires et al., 2016; Rai, 2002; Sharma and Dietz, 2006; Stewart and Larher, 1980; Warren et al. 2011; 2012). We found an overall upregulation of free amino acids in High and Low-pines, which also showed the most contrasted profiles (Fig. 5b; Table A.4), suggest that trees growing at the tree line and the rear-edge of the population distribution need to upregulate several metabolic pathways to cope with suboptimal environments.

Proline is a well-recognized multifunctional biomarker amino acid that plays critical roles in cellular redox balance (NADP^+^/NADPH ratio) (Szabados and Savouré, 2010), and tends to rapidly accumulate in plants under water stress (Alexieva et al., 2001; Bates et al., 1973; Dien et al., 2019; Singh et al., 1972; Yamada et al., 2005) as a response to the oxidative stress caused by drought (Alexieva et al., 2001; Moran et al., 1994; Munné-Bosch and Peñuelas, 2004). Pines at the tree line had the largest relative abundance of proline (Fig. 5b; Table A.4). Proline at high altitudes could play a prevalent function to cope with non-optimal environmental conditions that also induce oxidative stress. In fact, a strong correlation between ultraviolet-B (UV-B) incidence, which tends to be higher at high altitudes (Körner, 2007), and proline accumulation has been found in poplars (Ren et al., 2006). This correlation has also been observed in wheat and pea plants, where UV-B caused greater oxidation damage than drought (Alexieva et al., 2001). Supporting the potential antioxidant role of proline role in High-pines, those also presented higher relative abundance of tyrosine (Fig. 5b; Table A.4), which is a key precursor of phenylpropanoids. Phenylpropanoids are natural antioxidants (Korkina, 2007) that accumulate in plants under different pressures, such as UV irradiation, pathogens, high temperature, among others (Dixon and Paiva, 1995). In addition, High-pines showed higher relative abundances of glycin-betaine (GB) and its precursor glycine (Fig. 5b; Table A.4). GB has been reported as an important regulator in protection against oxidative plant stress caused by abiotic factors (Fariduddin et al., 2013). While GB has not been proven to possess direct antioxidant functions, it still activates ROS defensive pathways helping to cope with oxidative stress (Fariduddin et al., 2013; Ma et al., 2006; Smirnoff and Cumbes, 1989).

Similarly to carbohydrates and amino acids, pines growing at distinct altitudes also presented different profiles of organic acids, especially between the two lowest and the two highest altitudes (Fig. 5c; Table A.4). In general, several organic acids were found upregulated in Low pines (Fig. 5c; Table A.4). The tricarboxylic acid (TCA) cycle is central and a mechanistic understanding of the impacts of abiotic stress, such as drought, on their regulation and dynamics is still deficiently described (Fàbregas and Fernie, 2019). We did not find clear increases of fumaric, malic, oxaloacetic, aconitic, and succinic acids in Low-pines, which have been commonly associated with drought stress (Ashrafi et al., 2018; Fernández de Simón et al., 2022; Sardans et al., 2020; Urano et al., 2009). Citric acid (CA), on the other hand, was found upregulated in Low-pines (Fig. 5c; Table A.4). CA has been one of the most studied metabolites in plants under stress (Sardans et al., 2020) given its central role in the TCA cycle and being a key precursor of many metabolic pathways (Tahjib-Ul-arif et al., 2021). CA has been associated with many benefits in terms of plant stress tolerance, such as the upregulation of antioxidants and osmoregulators, as well as serving as a substrate for different pathways that synthesize stress-protectant compounds (Tahjib-Ul-arif et al., 2021). Generally, upregulation of CA has been found in most studies of plants under drought or heat stress (Sardans et al., 2020).

Several studies on exogenous application of CA to plants have generally found improved plant tolerance to drought (El-Tohamy et al., 2013; Tahjib-Ul-arif et al., 2021; Xie et al., 2022). Instead, other studies found that the response of CA to drought was dependent on the drought intensity and plant species (Ashrafi et al., 2018). Even though our results of CA align with most of the published research (Sardans et al., 2020), showing upregulation in pines growing in the most xeric area (Low-pines), the overall dynamics of the TCA cycle and the rapid turnover of specific metabolites could provide conflicting interpretations. Therefore, more holistic metabolomics approaches are needed to decipher the mechanistic temporal relationships between TCA compounds and stress, and to identify common and unique trends among different plants exposed to divergent environmental conditions and plant species.

We found a negative overall tendency of ascorbic acid from high to lower altitudes in *P. uncinata*, being the two lowest altitudes with higher relative abundances. (Fig. 5c; Table A.4). Although statistical inference did not detect ascorbic acid abundances changing significantly between altitudes, it is known to play critical roles as an antioxidant and radical scavenger in response to oxidative stress (Fàbregas and Fernie, 2019; Foyer and Noctor, 2011; Smirnoff and Wheeler, 2000) and most reports of plant water-limitation stress found upregulation (Jubany-Marí et al., 2010; Niu et al., 2013; Noman et al., 2015; Sun et al., 2020). However, glutathione (reduced form; GSH), the close partner of ascorbic acid in the ascorbate-glutathione cycle to detoxify hydrogen peroxide (H_2_O_2_) was clearly upregulated in Low-pines (Fig. 5c; Table A.4).

The glutathione and ascorbic acid profiles along the altitudinal gradient could be indicative of increased oxidative stress in Low-pines, in contrast to the intermediate locations. However, although glutathione has been typically associated with radical scavenger roles, similar to ascorbic acid (Foyer and Noctor, 2011; Noctor et al., 2012), research on glutathione synthesis, regulation and turnover indicates that its cellular dynamics in plants are more complex and not yet fully understood despite many oxidative stress processes are glutathione dependent (Noctor et al., 2012). In fact, the redox potential of glutathione is known to be influenced by the ratio of GSH to oxidized glutathione (GSSG) instead of only its absolute concentration (Mullineaux and Raush, 2005). It is essential to understand the turnover in the abundance of ascorbic acid, and the GSH and GSSG forms of glutathione in plants responding to specific stressors or environmental conditions. Plants do not rely on a single strategy to cope with the environment but activate an arsenal of mechanisms that do not necessarily follow identical temporal response patterns (Fàbregas and Fernie, 2019), which at the same time, can interact in redox cellular regulation (Noctor et al., 2012).

Glycolic and glyoxylic acids, both found in higher relative abundance in Low-pines (Fig. 5c; Table A.4), are key compounds of the photorespiration process to transform 2-phosphoglycolate (PG), a toxic metabolite resulting from O_2_ fixation by the enzyme ribulose 1,5-bisphosphate carboxylase-oxygenase (RuBisCO) when it accumulates in plant cells (Dusenge et al., 2019; Ogren, 1984; Zelitch et al., 2009). Low-pines, although not measured in this study, could present higher photorespiration rates compared to pines growing at higher altitudes, which grow under lower temperatures (Fig. 3a). The oxygenase activity of RuBisCO could be enhanced by the higher temperature at low altitudes (Jordan and Ogren, 1984). In addition, the observed increases in oxalic acid in Low-pines – despite ongoing debates about its biosynthesis pathways (Yu et al., 2010) - could be a byproduct of glycolate in the photorespiration process (Franceschi and Nakata, 2005). Oxalic acid has also been described as a natural antioxidant (Kayashima and Katayama, 2002) and an active osmotic compound (Li et al., 2022).

The shikimate pathway represents the starting point for several plant secondary metabolic processes that lead to diverse compounds (plant hormones, phenylpropanoids, flavonoids, alkaloids,…) with important roles in the ecophysiological response to the environment, especially under situations of abiotic (Guo et al., 2014; Herrmann, 1995; Jan et al., 2021) or biotic stress (Alvarez et al., 2016; Jan et al., 2021; Kumar et al., 2023). We found a gradual upregulation of shikimic acid from high to lower altitudes (Fig. 5c; Table A.4) which may suggest a high demand for secondary plant metabolism products. However, eco-metabolomics studies have not revealed yet a clear pattern in the relative abundance of the shikimic acid in plants under drought stress. Nonetheless, the literature indicates that several products of shikimic acid are generally found upregulated under water stress conditions (Sardans et al., 2020). Those inconclusive results may stem from varying rates of shikimic acid turnover among different plant species and environmental conditions. Given the ambiguous dynamics of shikimic acid found between plants under drought or heat stress, integrating metabolomics with transcriptomics and proteomics analyses is essential to begin clarifying its dynamics in plants growing in different environmental conditions.

### 4.3. Secondary metabolites shift of *Pinus uncinata* along altitudinal gradients

Plant secondary metabolism (PSM) includes phenolics, alkaloids, and steroids, terpenes, among others, and play critical functions in plant acclimation and adaptation, especially in response to environmental stressors (Bennett and Wallsgrove, 1994; Bourgaud et al., 2001). Single compounds from the PSM can have diverse functions, such as antifungal, antiviral, antibiotic, and hold anti-feeding properties. PSM compounds are well known to play an important antioxidant role against ROS, which include superoxide ion (O_2_^·−^), hydrogen peroxide (H_2_O_2_), hydroxyl radical (OH·), and singlet oxygen (^1^O_2_) (Larson, 1988; Mittler, 2002). In general, our results of the PSM suggest an overall increase of antioxidant activity in pines growing on both extremes of the altitudinal gradient (Fig. 6a, b; Table A.4). Most of the identified flavonoids have proven to play critical roles against a wide range of stresses, especially acting as antioxidants (Blokhina et al., 2003; Pietta, 2000; Rice-Evans et al., 1997). Following High and Low-pines, several phenolic compounds were also found upregulated in Int.1-pines suggesting thus an increased antioxidant function if compared to Int.2-pines which exhibited the lowest antioxidant activity (Fig. 6a, b; Table A.4). This result suggests that Int.2-pines probably thrive in a more optimal zone within their metabolome flexibility and can allocate more carbon to other functions such as storage, growth, and reproduction, rather than to defensive mechanisms. In accordance with the bibliography (Adams et al., 2017; Allen et al., 2010; Camarero et al., 2021; Gea-Izquierdo et al., 2021; McDowell et al., 2011; Pérez-Luque et al., 2021), the increased abundance of osmolytes and antioxidants found in Low-pines could be attributed to drier and warmer conditions if compared to pines growing at more optimal environments (Fig. 1). Previous research on the same *P. uncinata* population reported foliar injuries in High-pines that were directly associated with increased tropospheric O_3_ exposure (Díaz-de-Quijano et al., 2016). Although the O_3_ exposure, together with the wetter conditions at high altitudes, could explain the enhanced foliar antioxidant capacity in High trees, our metabolomics results cannot definitely confirm whether the increased oxidative stress found in High-pines is exclusively induced by O_3_ exposure. In fact, UV radiation, chilling; variables that tend to be more intense at higher altitudes (Körner, 2007), and O_3_ exposure, have proven to increase cellular ROS (Gill and Tuteja, 2010; Kovtun et al., 2000; Prasad et al., 1994), and plants respond upregulating secondary metabolic pathways for the synthesis of antioxidants such as flavonoids (Gill and Tuteja, 2010; Graf, 1992). Nevertheless, multifactorial studies in controlled chambers combining different levels of UV radiation, O_3_ exposure, and temperature are necessary to better understand the PSM reconfigurations in *P. uncinata* under a combination of factors producing oxidative stress.

The foliar relative abundance of over 50% of the identified phenolic compounds significantly correlated with altitude (Fig. 6c) indicating a gradual shift in response to environmental conditions changes (Fig. 3). However, phenolic compounds can also be upregulated in plants under a variety of conditions producing oxidative stress such as drought, warming, high-UV exposure, among other factors (Demidchik, 2015) which do not necessarily correlate with altitude. We observed an overall upregulation of the several phenolic compounds in pines growing at the edges of the altitudinal distribution of the population. This suggests that contrasted environments can still trigger similar oxidative stress responses in plants. Unlike correlation analyses (Fig. 6c), those trends can be more difficult to detect using multivariate approximations as a single compound could be upregulated in both Low and High-pines if compared to intermediate pines (Fig. 6a). Still, multivariate approximations such as PCA provide information of the weight of different metabolites relative to the environment. Therefore, eco-metabolomics studies performed along environmental gradients can provide important clues on which plant metabolic pathways are prioritized under the environmental conditions defined across altitudinal gradients (Fig. 7). Obtaining data on specific metabolic responses in different environments, including both primary and secondary metabolism, can help to identify more suitable biomarkers for specific environmental conditions.

**Figure 7.**
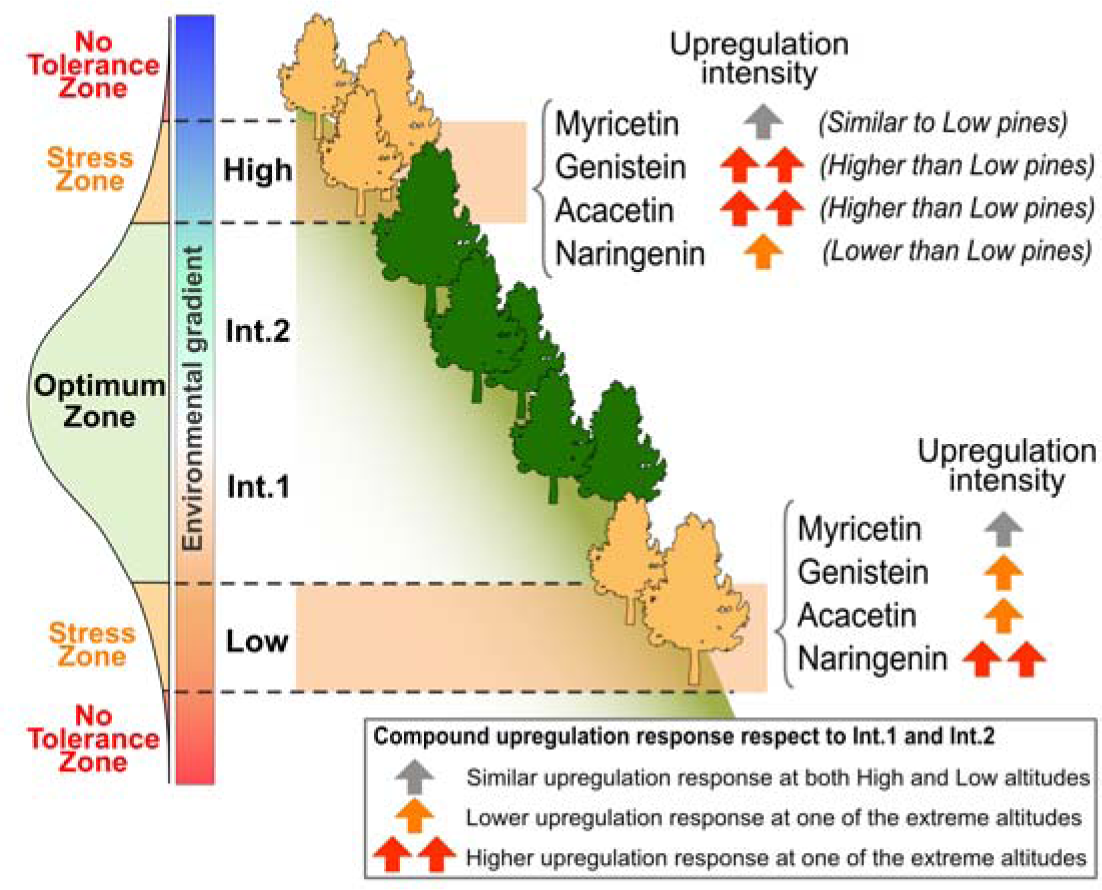
Plant differential upregulation intensity to oxidative stress. Conceptual illustration of the plant differential upregulation intensity of phenolic compounds in response to oxidative stress produced by contrasting environments. Although both High and Low-pines upregulate several phenolic compounds, its intensity diverge in function of the environment they grow. This shows a preference of specific antioxidants to cope with the oxidative stress produced by the particular environment of the lowest and highest altitudes.

## 5. Conclusions and final remarks

Eco-metabolomics approaches along environmental gradients offer a valuable means to explore and proxy the overall metabolome flexibility of natural plant populations. The chemical phenotypic flexibility of a population, therefore, is understood as a proxy of the overall flexibility of genotypes within the studied population. As hypothesized, we found evidence of increased metabolite activity in pines growing at both edges of the altitudinal distribution of the population. Trees at the highest and lowest altitudes exhibited large shifts in their overall metabolomes (Fig. 4), accompanied by changes in sugars, amino acids, and antioxidants commonly associated with oxidative stress (Figs. 5, 6). However, it is important to note that the distribution range of plant populations can also be influenced by other abiotic and biotic factors beyond climate, such as competition, physical barriers, and soil structure (Fig. 2). These factors may complicate deciphering the metabolomic flexibility of natural plant populations. Therefore, obtaining more accurate measurements of a plant population’s metabolome flexibility may require complementing field measurements with chamber and common garden experiments which expose plants to controlled environmental gradients, aiding in elucidating the metabolic mechanisms underlying acclimation versus genetic responses. Establishing reliable indices of chemical phenotypic flexibility for plant populations and species is necessary for assessing the ecosystem structure and function under the current scenarios of global change and thereby providing critical insights into the future of plant populations.

In addition to the challenges presented by the rapid pace of climate change, our findings suggest that upward migration in response to global warming may still not be feasible for some plant populations due to the presence or the increased intensity of other environmental variables that cause oxidative stress. Increased UV and/or O_3_ exposure have the potential to induce oxidative stress, reducing individual fitness and the capacity of plant individuals to thrive beyond those boundaries. Although the increased oxidative foliar damage and forest decline in high trees of the studied *P. uncinata* population has previously been directly associated with an enhanced tropospheric O_3_ exposure, metabolomics analyses alone cannot definitively conclude whether O_3_ is the main factor leading to increased foliar oxidative stress at high altitudes. To address this, stress-specific biomarkers or metabolic signatures need to be identified and integrated into metabolomics plant libraries. In this way, our study suggests that eco-metabolomics measurements of plants along their altitudinal distribution have the potential to identify key and more meaningful compounds that can serve as biomarkers, directly related to specific environmental conditions defined by particular variables (water availability, temperature,…) (Fig. 7).

To conclude, the result of our study underscores the significance of a robust experimental design in metabolomic studies conducted in natural conditions as plants respond to a combination of environmental variables that vary along environmental gradients.

## Author contributions

**Albert Rivas-Ubach:** Conceptualization, Data curation, Formal analysis; Funding acquisition, Project Administration, Investigation, Methodology, Writing - original draft, **Ismael Aranda:** Conceptualization, Investigation, Writing - review & editing, **Jordi Sardans:** Supervision, Investigation, Funding acquisition, Writing - review & editing, **Yina Liu:** Investigation, Writing - review & editing, **María Díaz de Quijano:** Investigation, Writing - review & editing, **Ljiljana Paša-Tolić:** Resources, Methodology, Funding acquisition, Writing - review & editing, **Michal Oravec:** Resources, Methodology, Writing - review & editing, **Otmar Urban:** Resources, Methodology, Funding acquisition, Investigation, Writing - review & editing, **Josep Peñuelas:** Supervision, Investigation, Writing - review & editing, Funding acquisition, Project administration

## Acknowledgements

This research was supported by a research fellowship (JAE) from the CSIC, the Autonomous Agency of National Parks of Spain (OAPN) project 2948/2022 (MetForDec), the Spanish Government grants PID2022-142746NB-I00, PID2020-115770RB-I, PID2022-140808NB-I00, and TED2021-132627B–I00 funded by MCIN, AEI/10.13039/ 501100011033 European Union Next Generation EU/PRTR, the Fundación Ramón Areces grant CIVP20A6621, and the Catalan Government grants SGR 2021–01333 and AGAUR2023 CLIMA 00118. MO and OU were supported by the project of the Ministry of Education, Youth and Sports of the Czech Republic (AdAgriF; CZ.02.01.01/00/22_008/0004635). A portion of this research was performed at the Environmental Molecular Sciences Laboratory (EMSL), a DOE Office of Science User Facility sponsored by the Office of Biological and Environmental Research at Pacific Northwest National Laboratory.

## Notes

### Competing Interest Statement

The authors have declared no competing interest.

