## Appendix A for "An outline on the chemical phenotype flexibility of forest species: an eco-metabolomics study of Pinus uncinata along an altitudinal gradient"

Metabolite extraction for LC-MS

Metabolites were extracted with methanol:water (80:20) for a broad recovery of polar and semi-polar metabolites. Two separate sets of 2 mL glass vials were labeled (referred here as set A and set B). Set A was to perform metabolite extractions, while set B was to keep the extracts for subsequent LC-MS analyses. Briefly, for each sample, 100 mg of lyophilized powder was weighted into the corresponding vial of set A, and 1 mL of methanol:water solution was subsequently added. Vials, with polytetrafluoroethylene (PTFE) caps, were vortexed for 15 min, sonicated for 3 min at room temperature (22 °C), and centrifuged at 10, 000 × g for 5 min. Supernatants (0.6 mL) were collected and transferred to the corresponding vials of set B. By adding 0.6 mL of fresh methanol:water solution to set A, the previous steps were repeated for an additional extraction from each sample and 0.6 mL were thus pooled with the first extraction of the set of vials B. All extracts were individually filtered with 0.22 µm syringe microfilters and introduced into newly labeled high-performance liquid chromatograph (HPLC) glass vials.

LC-MS analyses

LCMS analyses were conducted on a Dionex Ultimate 3000 HPLC (Thermo Fisher Scientific, USA) coupled to a high-resolution mass spectrometer (HRMS) Orbitrap XL equipped with a HESI II heated electrospray ionization source. The C18 column (Hypersil GOLD, 150 × 2.1 mm, 3 µm particle size; Thermo Fisher Scientific, [Waltham, USA](http://en.wikipedia.org/wiki/Waltham,_Massachusetts)) was maintained at 30 °C during the chromatography. The autosampler temperature was maintained at 8 °C. The mobile phases consisted of 0.1% acetic acid (A) and acetonitrile (B). Both phases A and B were filtered and degassed for 10 min in an ultrasonic bath prior to use. Chromatography was performed at a constant flow rate of 0.3 mL min^-1^ with 5 µL injections. The elution gradient began at 90% A (10% B) and was held for 5 min. A was thus linearly decreased to 10% during the following 15 min (min 20), and the solvent proportion was maintained for another 5 min (min 25). The chromatographic gradient returned thus to the initial conditions (10% B and 90% A), and the column was washed and equilibrated for 10 additional minutes (min 35) before the analysis of the next sample.

The HRMS operated in FTMS (Fourier transform mass spectrometry) full-scan mode at a resolving power of 60, 000. Full scan spectra were acquired over a mass range of 70–1000 m/z in both positive and negative modes. The sensitivity and resolution of the HRMS were regularly monitored by injecting a mixture of phenolic compounds (negative mode) and caffeine (positive mode).

LC-MS dataset filtering and normalization

The main aim of eco-metabolomics studies is to understand the main overall metabolome shifts and trends that organisms exhibit under particular conditions (e.g. pressure under biotic or abiotic stressors). To achieve this, a large number of biological replicates is typically analyzed to infer the overall effects of biotic or abiotic factors on the studied population. Commonly, in ecological studies, our primary aim is to reveal overall trends in populations, communities and ecosystems. For that reason, in order to achieve this global perspective, it is convenient to manage sporadic and non-representative metabolomic variables effectively. Those variables can arise from two primary sources of variation: i) mass spectrometry instruments and RAW data processing, and ii) the natural metabolomic plasticity of organisms. From the instrumental and bioinformatic perspective, mass spectrometry instruments generate a significant amount of background, and some signals may still be detected as “metabolic features” after data processing in MZmine or other software. Peak spikes are also a common source of non-representative features in some samples. In MZmine, all those features can originate from various data processing steps, especially during ion detection, ion chromatograms deconvolution, chromatogram aligner, and gap-filling. From a metabolic plasticity point of view, non-representative features can still be genuine and not artifacts resulting from poorly resolved chromatograms or MZmine algorithms, given the extensive plasticity exhibited by organisms, especially plants. Yet, when considering an ecological perspective that emphasized a broader view, individualities (e.g. presence of few metabolic features within a small number of individuals within a large population) cannot be considered as representative of the population and commonly do not provide additional insights from a broader perspective. Although, paying attention to some idiosyncrasies in ecophysiological studies may be interesting, our study focuses on overall trends along altitudinal gradients and the LC-MS datasets obtained in negative and positive ionization modes were filtered independently at *cell* level to reduce the number of noisy and non-representative features (e.g., signals detected in a one or two samples out of 40). A *cell* of the dataset corresponds to all replicate samples within the same experimental group. In our case, given that our study only contains a single categorical factor, altitude, it has 4 different *cells*: Low, Int.1, Int.2, and High.

Dataset filtering was performed through 3 primary steps:

(1) *Minimum data*. Metabolite features with data (peak areas >0) in less than 70% of the replicates (6 samples or less out of 10) of the 4 *cells* were removed from the dataset. Only those features with at least 1 *cell* containing data in 70% or more of its replicates were maintained.
(2) *Replicate identified features simplification and data merging.* Instrument measurement error (typically <5 ppm) and spikes on chromatographic peaks may provide separate features with slight m/z and/or RT values. Those features may fall within the tolerance error for both exact mass and RT parameters during library compound matching, providing thus 2 or more features with the same identification. Therefore, all identified ions within the same ionization mode assigned to the same molecular compound were summed to obtain a single variable per identified compound. Subsequently, the datasets originated from positive and negative ionization modes were merged into one. Then, if the same compound identity has been found in both positive and negative ionization datasets, it is merged into a single variable adding up both values weighting their relative abundances accordingly to avoid overrepresentation of one of the ionization modes.
(3) *Total chromatogram intensity normalization and scaling.* The values of a sample for each detected feature were normalized by the total chromatogram area of the sample. Then, the metabolite features (continuous variables) were individually *mean-centered* and *auto-scaled* (divided by the standard deviation of the variable).

Multivariate outlier tests

Samples were submitted to multivariate outlier tests based on the Mahalanobis distance for potential outlier detection. The results of each test were shown through principal component analyses (PCA). The tests were performed with the *pca.outlier* function of the “mt” package of R. When considering all altitudes together at a confidence level of 95%, two High-pines (Replicates 1 and 2) had slightly larger Mahalanobis distance values (2.514 and 2.465, respectively) than the established cutoff of 2.448 (Fig. A.1; Table A.3). This suggests that replicates 1 and 2 are multivariate outliers. However, the analysis using the package default confidence level (0.975), no outliers were identified. When analyzing each altitude separately, no multivariate outliers were detected at a confidence of 0.95 for any of the altitudes (Fig. A.1; Table A.3). Therefore, since no extreme outliers were detected when analyzing all altitudes together and no outliers were identified within each group, all samples were thus included in the subsequent statistical analyses.

**Fig. Appendix A.1.** Principal component analyses (PCAs) with Mehalanobis distance thresholds shown across PC1 and PC2 axes. The analyses were computed at a confidence level of 0.95 for all samples together (a), Low pines (b), Int.1 pines (c), Int.2 pines (d), and High pines (e). High pines at a confidence of 0.975 (f) and High and Int.2 pines together at a confidence level of 0.95 (g) were also evaluated. Replicate 5 of High pines is pointed by a black arrow. Samples outside the red ellipse is considered an outlier sample for the confidence level established.
