## Supplementary material for "An outline on the chemical phenotype flexibility of forest species: an eco-metabolomics study of Pinus uncinata along an altitudinal gradient": Figure A.1

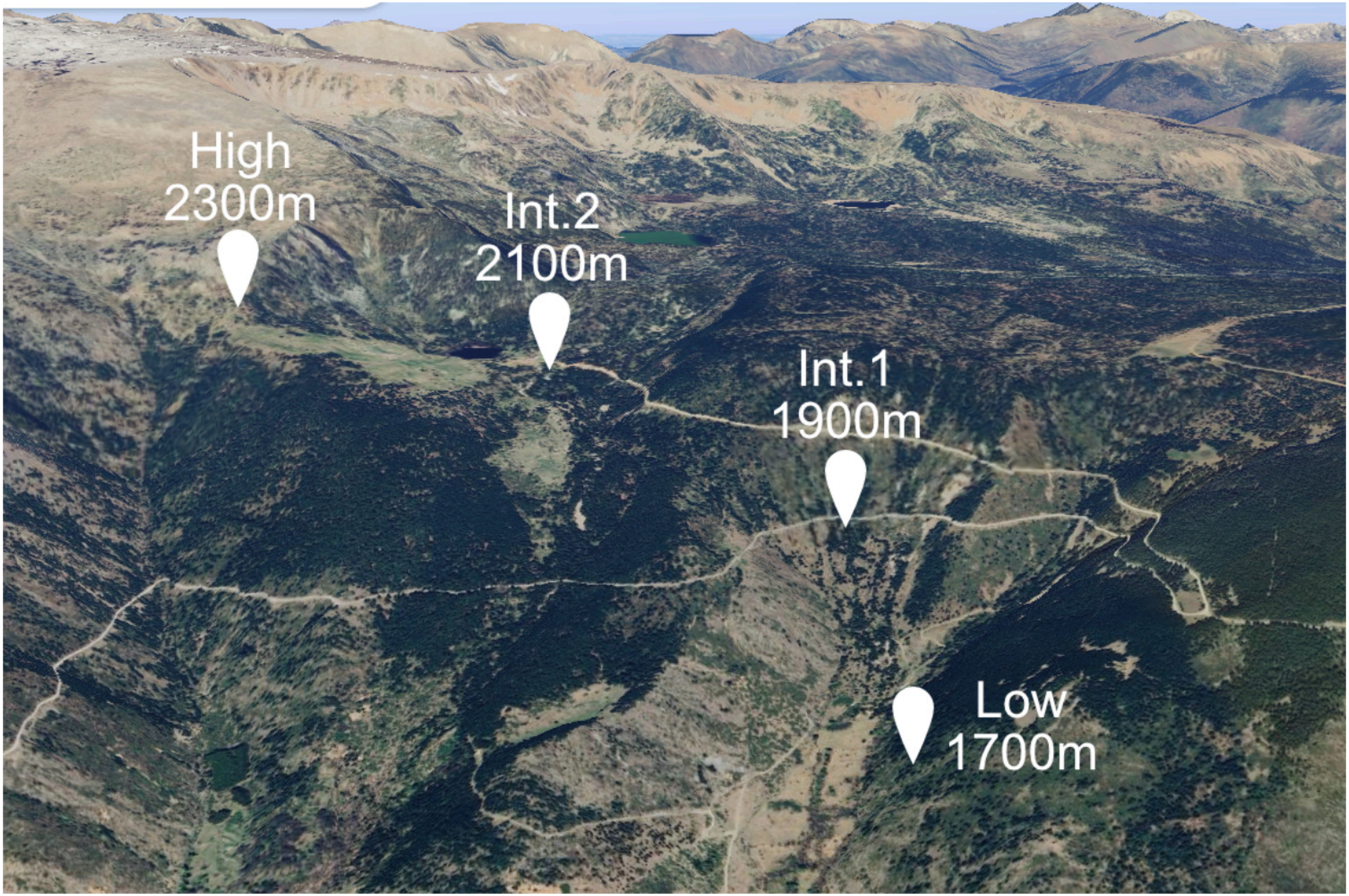


**Figure A.1** Location of the four different altitudes studied along the altitudinal gradient of the population of *Pinus uncinata* in the valley of Meranges (Catalonia, Spain).
