## Supplementary material for "An outline on the chemical phenotype flexibility of forest species: an eco-metabolomics study of Pinus uncinata along an altitudinal gradient": Figure A.2

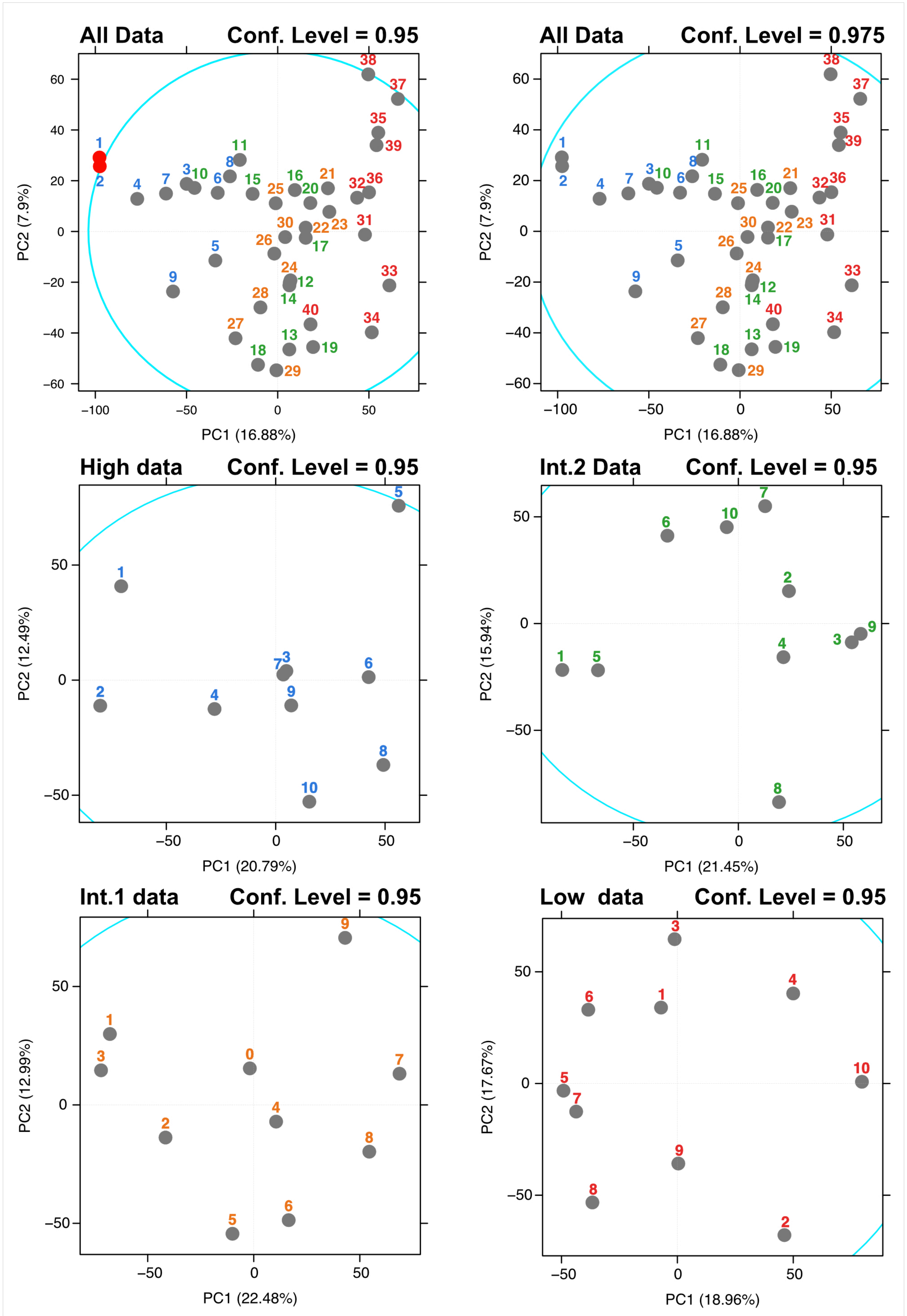


**Figure A.1** Principal Component (PC)1 vs. PC2 of the Principal Component analyses (PCAs) for all metabolomics samples and for each of the altitudes. The Mahalanobis distance is represented in the plots with a light-blue ellipse, which measures the distance of each point from the center of the distribution. Samples plotted outside the Mahalanobis distance can be considered as multivariate outliers. The different confidence levels tested are indicated above each PCA.
