## Supplementary material for "An outline on the chemical phenotype flexibility of forest species: an eco-metabolomics study of Pinus uncinata along an altitudinal gradient": Figure A.3

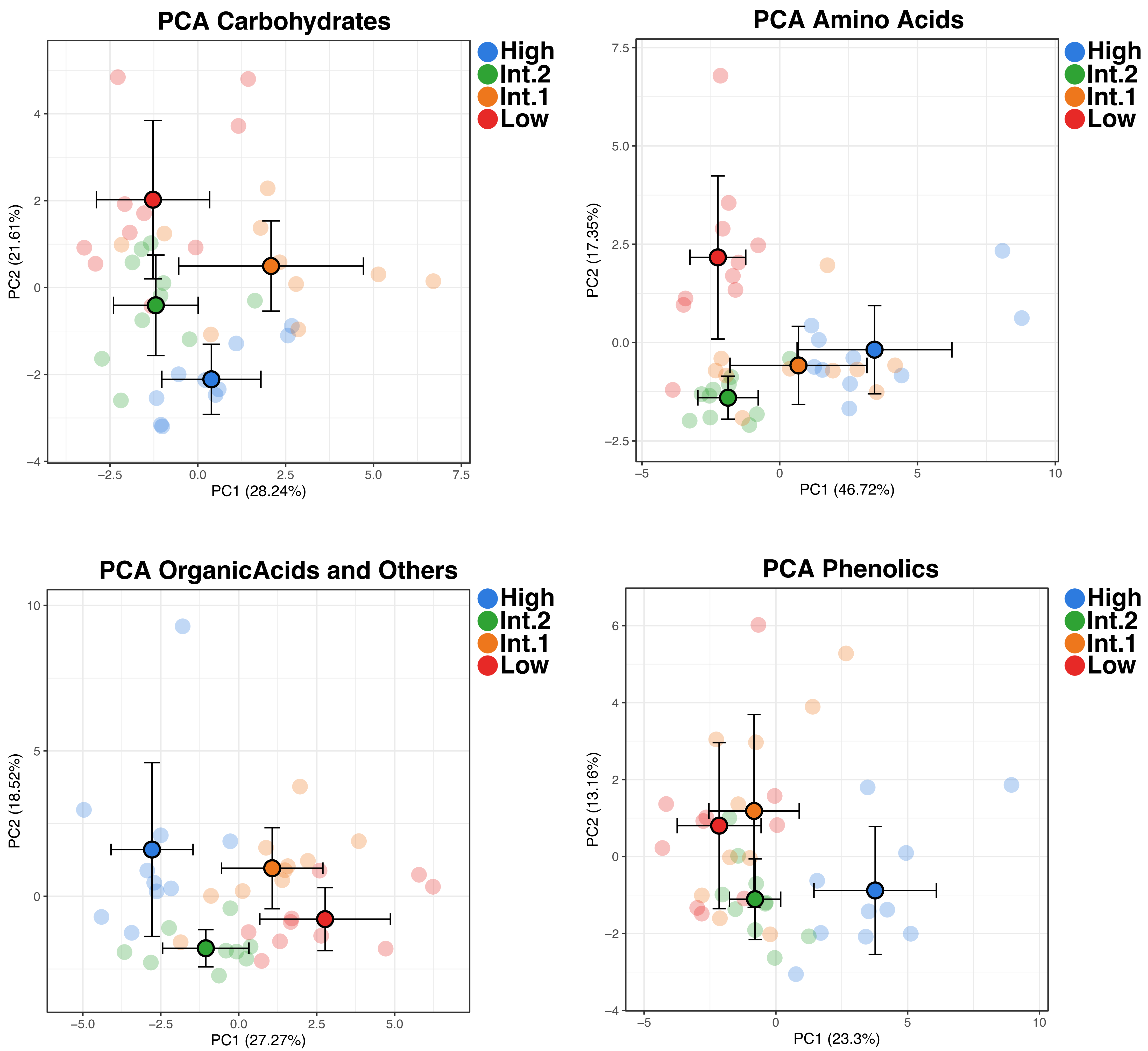


**Figure A.** Principal component (PC) 1 vs. PC2 of the principal component analysis (PCA) for carbohydrates, amino acids, organic acids and others, and phenolic compounds. Different altitudes are shown in different colors (Low in red; Int.1 in orange; Int.2 in green; High in blue). The samples across the 2D plane are represented with faded dots and the average value for each altitude is shown in deep colored dots with black outlines. Standard deviation bars for the averaged values of each altitudes are shown for both principal components.
