## Supplementary material for "An outline on the chemical phenotype flexibility of forest species: an eco-metabolomics study of Pinus uncinata along an altitudinal gradient": Figure A.4

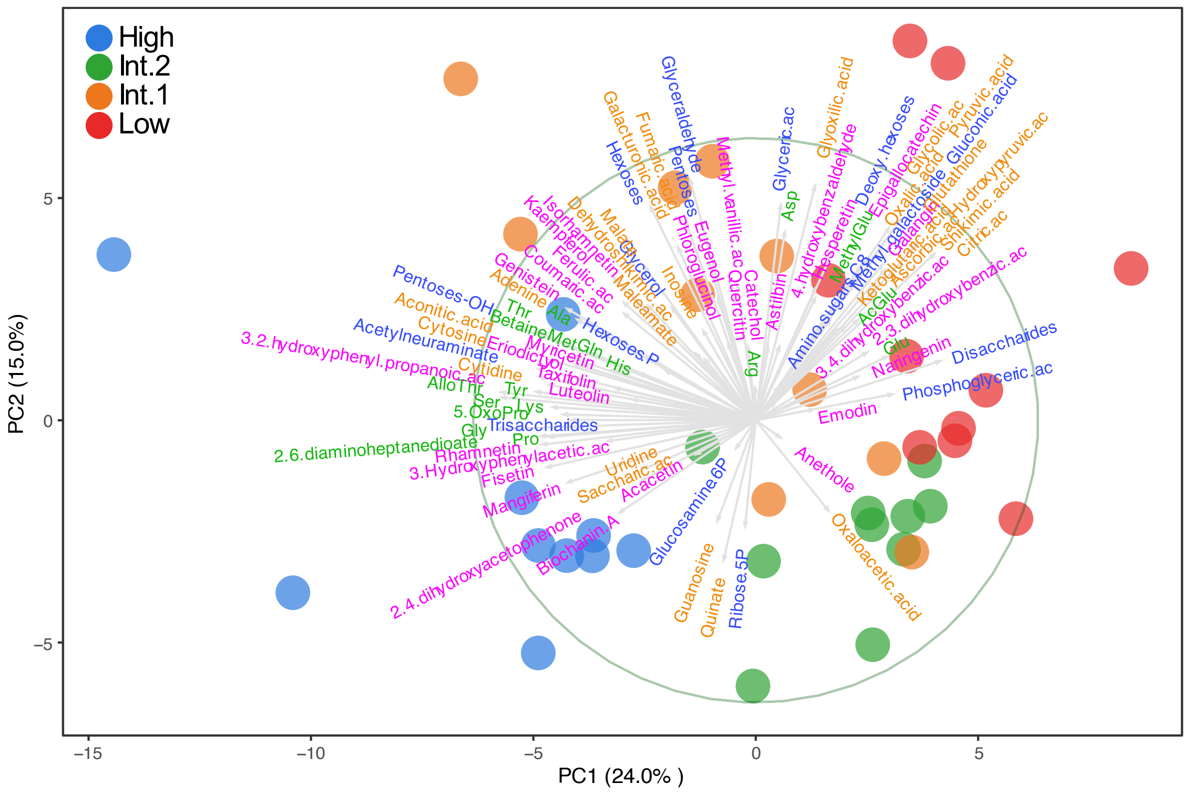


**Figure A.2**. Principal component (PC) 1 vs. PC2 (**a**) of the biplot of the principal component analysis (PCA) performed with the metabolomic compound classes (carbohydrates in blue; amino acids in green; phenolics in purple; organic acids and other compounds of interest in orange) identified in needles of *Pinus uncinata*. Pines from different altitudes are shown with different colors (Low in red; Int.1 in orange; Int.2 in green; High in blue).
