## Supplementary material for "An outline on the chemical phenotype flexibility of forest species: an eco-metabolomics study of Pinus uncinata along an altitudinal gradient": Figure A.5

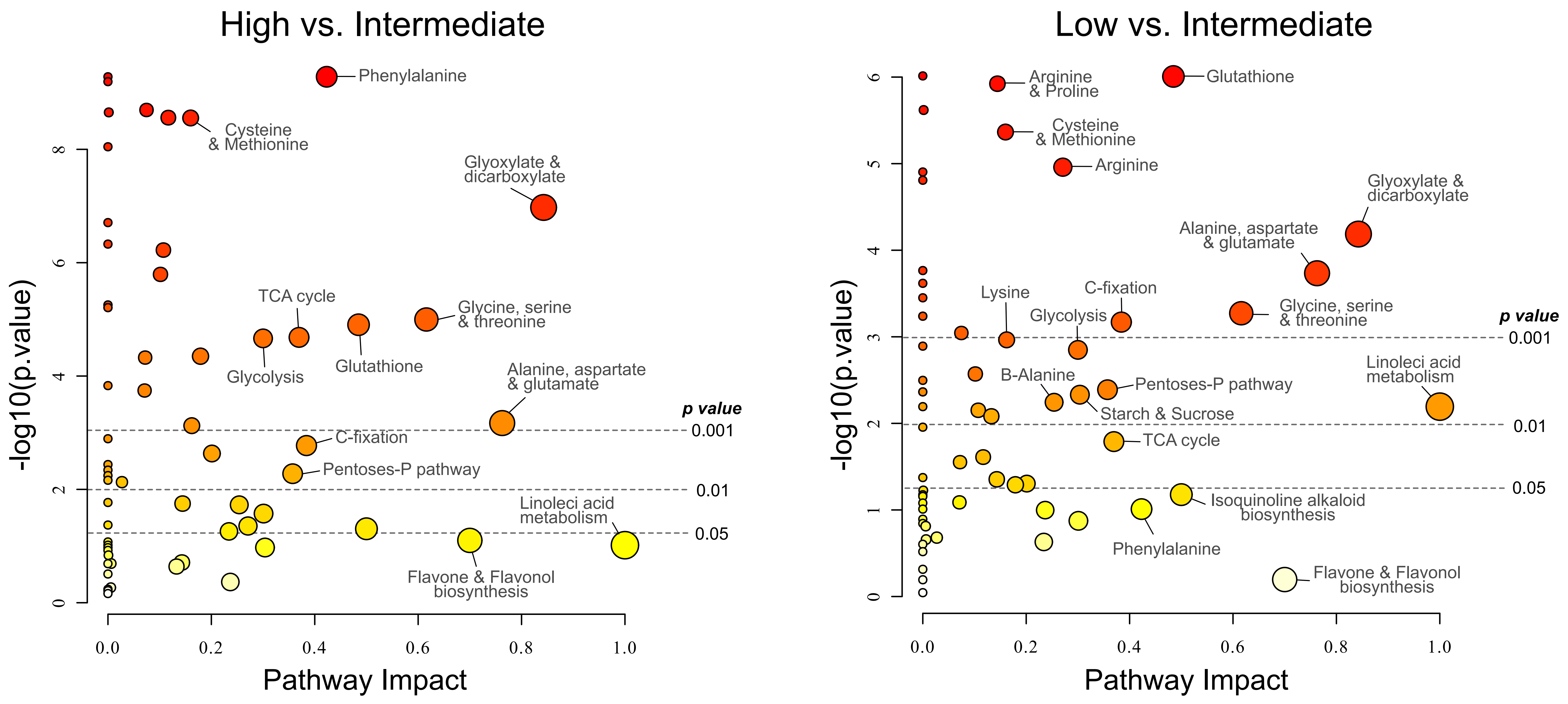


**Figure S1.** Pathway analysis plots of Low-pines vs. Intermediate-pines (Int.1 and Int.2), and High-pines vs. Intermediate-pines (Int.1 and Int.2) performed in Metaboanalyst 5.0 (Pang et al., 2021). Pathway impact (Xaxis) represents a combination of enrichment analysis and centrality of changing compounds within the pathway. Y-axis indicates the -log10(p value) of represented pathways. Absolute P value thresholds of 0.05, 0.01 and 0.001 are indicated with dashed lines. Absolute pathway P values are represented in different colors being red the most significant values (lower P value) and light-yellow, the least significant values (higher P value).

References.

Pang, Z., Chong, J., Zhou, G., De Lima Morais, D.A., Chang, L., Barrette, M., Gauthier, C., Jacques, P.É., Li, S., Xia, J., 2021. MetaboAnalyst 5.0: Narrowing the gap
